# Mapping the risks of China’s global coastal development to marine socio-ecological systems

**DOI:** 10.1101/2022.04.22.489174

**Authors:** B. Alexander Simmons, Nathalie Butt, Casey C. O’Hara, Rebecca Ray, Yaxiong Ma, Kevin P. Gallagher

## Abstract

Rapid coastal development continues to jeopardize the integrity of marine socio-ecological systems. China is now the largest bilateral creditor in the world, committing nearly half a trillion US dollars to overseas development finance since 2008. Meanwhile, there are growing concerns over the impacts of this boom in Chinese development finance on marine systems. Here, we quantify the risks of coastal development projects financed by China to marine biodiversity and coastal Indigenous communities. Ports present the greatest impact risks to marine systems, in terms of both magnitude and area at risk, with power plants, roads, and other facilities presenting relatively high localized risks. Risks are most prominent in Africa and the Caribbean, with coastal Indigenous communities in Western and Central Africa particularly vulnerable to the potential negative impacts of development. All projects present some risk to threatened marine species and potential critical habitats, but few present high risks to nearby marine protected areas. Most projects present additional risks to ecosystems that are already under increasing human pressures, but some are likely to introduce new risks to relatively intact ecosystems. “Bluing” future coastal development projects in China’s overseas development finance portfolio will require more social and environmental safeguards, higher standards for host-country impact assessments, and greater integration of land-sea risk mitigation and management approaches.

## Introduction

The state of the world’s oceans is in rapid decline from the growing impacts of climate, land, and ocean-based threats^1^. The intensity of these anthropogenic impacts is often greatest in coastal waters^1, 2^, which are one of the top priority areas for marine biodiversity conservation^3^. The continual growth of infrastructure and pollution associated with coastal development threatens the structure and function of marine socio-ecological systems around the world^4–6^. Urbanization, sedimentation, and nutrient input represent some of the most common causes of marine regime shifts, including ecosystem transitions and collapse, hypoxia, and eutrophication^7^, which will ultimately impact more than 37% of the human population living in coastal communities that rely on the ecosystem services of these aquatic habitats^8^. With coastal populations expected to continue increasing over the next three decades^9^, “bluing” coastal development activities remains one of the major priorities for achieving several UN Sustainable Development Goals, such as ending poverty (Goal 1), economic growth (Goal 8), sustainable cities and communities (Goal 11), and conserving life on land and sea (Goals 14, 15)^10, 11^.

The last decade has brought a boom in overseas financing for large-scale development, with Chinese development finance institutions (DFIs) emerging as the largest bilateral creditor of these development projects^12, 13^. The China Development Bank (CDB) and Export-Import Bank of China (CHEXIM), alone, have lent more than US$460 billion to low- and middle-income countries since 2008 for the development of roads, power plants, ports, and other infrastructure^14^. It is estimated that China is also the largest bilateral funder of coastal infrastructure overseas, surpassing Japan and multilateral DFIs, such as the Asian Development Bank and Inter-American Development Bank^15^. Furthermore, while other major DFIs are greatly reducing their financial support for infrastructure projects^16^, Chinese DFIs have risen to become the major source of infrastructure finance globally. Coastal infrastructure, in particular—including ports and the roads that connect those ports to inland highway and rail networks—have risen to comprise a major element of China’s recent foreign economic diplomacy^17, 18^. With China’s ascent as a global economic powerhouse came the development of the Belt and Road Initiative (BRI) in 2013—a long-term, multi-billion-dollar, transcontinental infrastructure investment program aimed at strengthening cooperation, trade, and connectivity between China and countries with a shared vision of economic growth and development^17^. The BRI, which includes the terrestrial-based “Silk Road Economic Belt” and the marine-based “21^st^ Century Maritime Silk Road,” has enormous potential to propel economic prosperity in emerging markets^18, 19^, yet there remain growing concerns over the potential deleterious impacts of this initiative on the environment and local and Indigenous communities^20, 21^.

In a recent study^22^, researchers investigated the risks implicit in China’s overseas development finance portfolio to terrestrial biodiversity and Indigenous lands and found these risks to generally be greater than comparable projects financed by the World Bank, particularly in the energy sector. Yet while some studies have discussed the potential implications of increased shipping traffic along the Maritime Silk Road on marine systems^23, 24^ or examined the proximity of BRI-affiliated ports to marine ecoregions and threatened species^25^, there remains no comprehensive, quantitative evaluation of the risks to marine socio-ecological systems across China’s diverse portfolio of overseas development finance. This gap is exceptionally concerning given that more than 27 million coastal Indigenous peoples around the world are reliant upon a healthy and functioning marine ecosystem for food and income^26^. Moreover, the scarcity of high-precision, georeferenced data on overseas development finance has impeded scholars’ ability to track any bilateral or multilateral development banks’ financing of global coastal development.

Here, we investigated the extent to which coastal development projects financed by China’s major DFIs present both direct and indirect risks of adverse impacts on marine ecosystems and the coastal Indigenous communities reliant upon them. We identified 114 DFI projects in 39 countries financed between 2008 and 2019 that may negatively impact marine socio-ecological systems, including ports, power plants, roads, and other facilities (Fig. 1a), representing one-fourth of all Chinese DFI projects that have been geolocated with the highest precision, and worth nearly $65 billion in finance commitments (7% of all commitments) since 2008^14^. For each different type of project, we estimated the distribution of exposure risks (*R_E_*)—the likelihood that a given ocean cell may be exposed to a particular stressor—up to 100 km from the project site for 15 stressors, such as habitat loss, invasive species, and pollution from light, noise, and organic and inorganic chemicals. We then estimated the impact risks (*R_I_*)—the likelihood that the stressor(s) will lead to adverse impacts—based upon exposure risk(s) and the vulnerability of the present habitats and species to the particular stressor(s) (see Methods).

**Fig. 1.**
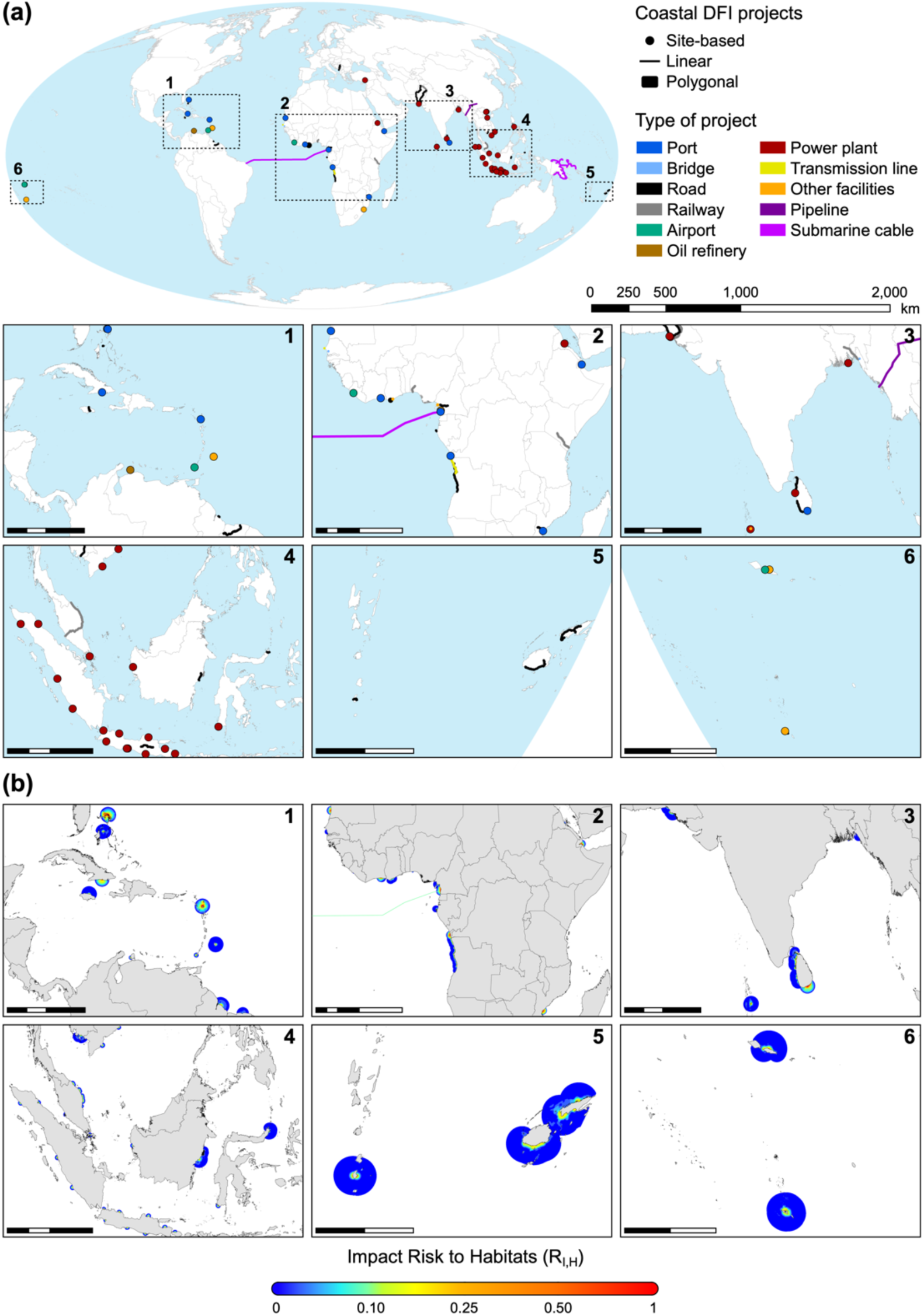
**(a)** Location of 114 projects financed by Chinese development finance institutions (DFI) during 2008–2019 that present some risk to coastal and/or marine systems. Lines and polygon boundaries thickened to enhance visibility. **(b)** Distribution of impact risks to near- and off-shore habitats (*R_I,H_*) from each DFI project site. See Supplementary Fig. 1 for larger maps of impact risks to each region.

We compared how these impact risks vary between different types of projects and how risk declines at increasing distances from the project site for 20 types of marine habitats based on their vulnerability to each stressor^27^. To identify where DFI projects pose new threats to habitats facing relatively few anthropogenic pressures, we also compared marine habitats’ impact risks with existing cumulative human impacts^1^. Finally, we estimated impact risks of DFI projects to the following socio-ecological features, which are exceptionally sensitive to the impacts of coastal development: threatened marine species, marine protected areas, areas likely to qualify as ‘critical habitat’, and areas that may be important resources for coastal Indigenous communities (referred to as ‘Indigenous-use seas’).

## Results

### Risks to marine habitats

Impact risks to marine habitats are most prominent surrounding island nations in the Caribbean, such as the Bahamas and Antigua and Barbuda, as well as coastal waters across Africa, most notably along Western and Central African coastlines (Fig. 1b). Impact risks in these regions tend to present greater magnitudes of risk, and over larger distances, compared with other regions. In the Bahamas, Angola, and Mozambique, more than 2000 km^2^ of marine habitats face high impact risks (*R_I,H_* > 0.25) (Extended Data Fig. 1a). Smaller impact risks (*R_I,H_* < 0.10) are more pervasive across countries; Angola, Fiji, Sri Lanka, and Indonesia, for example, have more than 50,000 km^2^ of their marine habitats facing low but non-negligible risks from nearby DFI projects (0 < *R_I,H_* < 0.10). Among recipient countries of Chinese overseas development finance, those with the smallest extent of risk include Myanmar (9.61 km^2^), Kenya (21.83 km^2^), and Liberia (306.51 km^2^), where impact risks never exceed *R_I,H_* = 0.10. Overall, some of the most at-risk habitats over larger distances include pelagic surface waters, shallow soft bottom ecosystems, and rocky reefs (Extended Data Fig. 2). Impact risks tend to be highest for most habitats 1–5 km from the project site, including shellfish/suspension reefs, salt marshes, mudflats, and rocky intertidal habitats.

Ports present the greatest impact risks to marine habitats around the world (*mean ± SD*: *R_I,H_* = 0.93 *±* 0.05), and these risks remain high even up to 30 km from the port (*R_I,H_* = 0.31 *±* 0.14) (Fig. 2). These ports are present in the Bahamas, Antigua and Barbuda, Cuba, Mauritania, Côte d’Ivoire, Cameroon, Angola, Mozambique, Djibouti, and Sri Lanka, and are a prominent driver of the regional hotspots of risk (Fig. 1). Most notably, the only fishing port included in this study— the Beira Fishing Port Rehabilitation project in Mozambique—presents the single greatest mean impact risk to marine habitats within 10 km of all projects considered in this study (*R_I,H_* = 0.80 *±* 0.05). Several other types of DFI projects present high impact risks within 1 km of the project site, such as power plants (*R_I,H_* = 0.40 *±* 0.07), bridges (*R_I,H_* = 0.38 *±* 0.09), roads (*R_I,H_* = 0.34 *±* 0.06), and other facilities (*R_I,H_* = 0.38 *±* 0.07). Impact risks decline rapidly, however, at increasing distances from the project site, with most types of projects presenting relatively low risks (*R_I,H_* < 0.10) beyond 10 km, on average (Fig. 2).

**Fig. 2.**
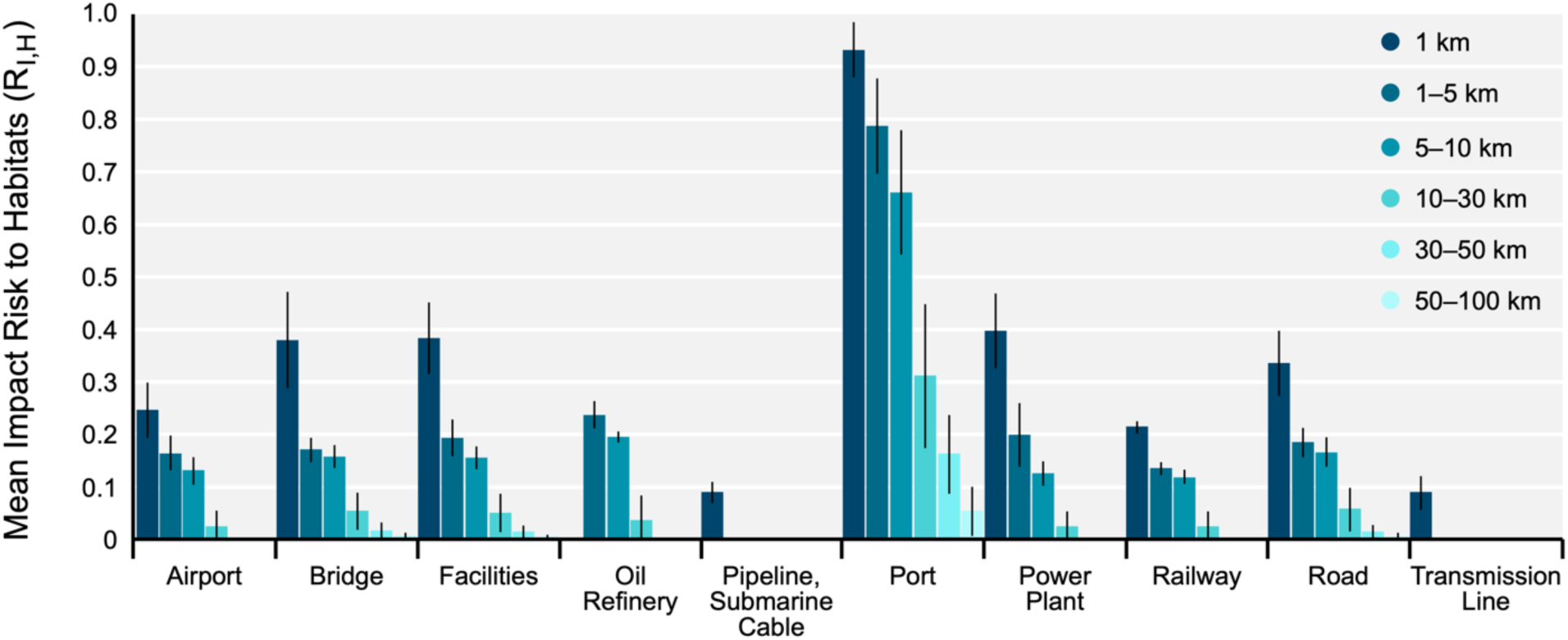
Mean impact risk to all habitats (*R_I,H_*) for different types of DFI projects at increasing distances from the project site. Error bars represent the standard deviation of the mean.

Overall, Chinese DFI projects present the greatest additive risks to marine habitats that are already experiencing high cumulative human impacts (CHI), with most projects characterized by very high mean CHI scores at time of financing—often several orders of magnitude higher than the respective global median CHI values for 2008 and 2013 (Fig. 3). Some projects, however, present risks to marine habitats that were previously under lower human pressures, such as the Doraleh and Damerjob Ports in Djibouti, the Kribi Port in Cameroon, and the North Abaco Port in the Bahamas. Furthermore, 79% of DFI projects present these additional impact risks to habitats that were experiencing increasing human impacts (ΔCHI > 0) from land- and ocean-based pressures since 2003 (Fig. 3). An additional 17 projects (15%) present risk to habitats where human impacts have been increasing or decreasing, on average, depending upon the distance from the project site. Only 7 projects (6%) consistently present impact risks to habitats that have previously seen a net reduction in human-induced impacts (ΔCHI < 0)—three of which are coal-fired power plants in Vietnam with moderately high impact risks up to 10 km from the plant (*R_I,H_* > 0.10) (Fig. 3).

**Fig. 3.**
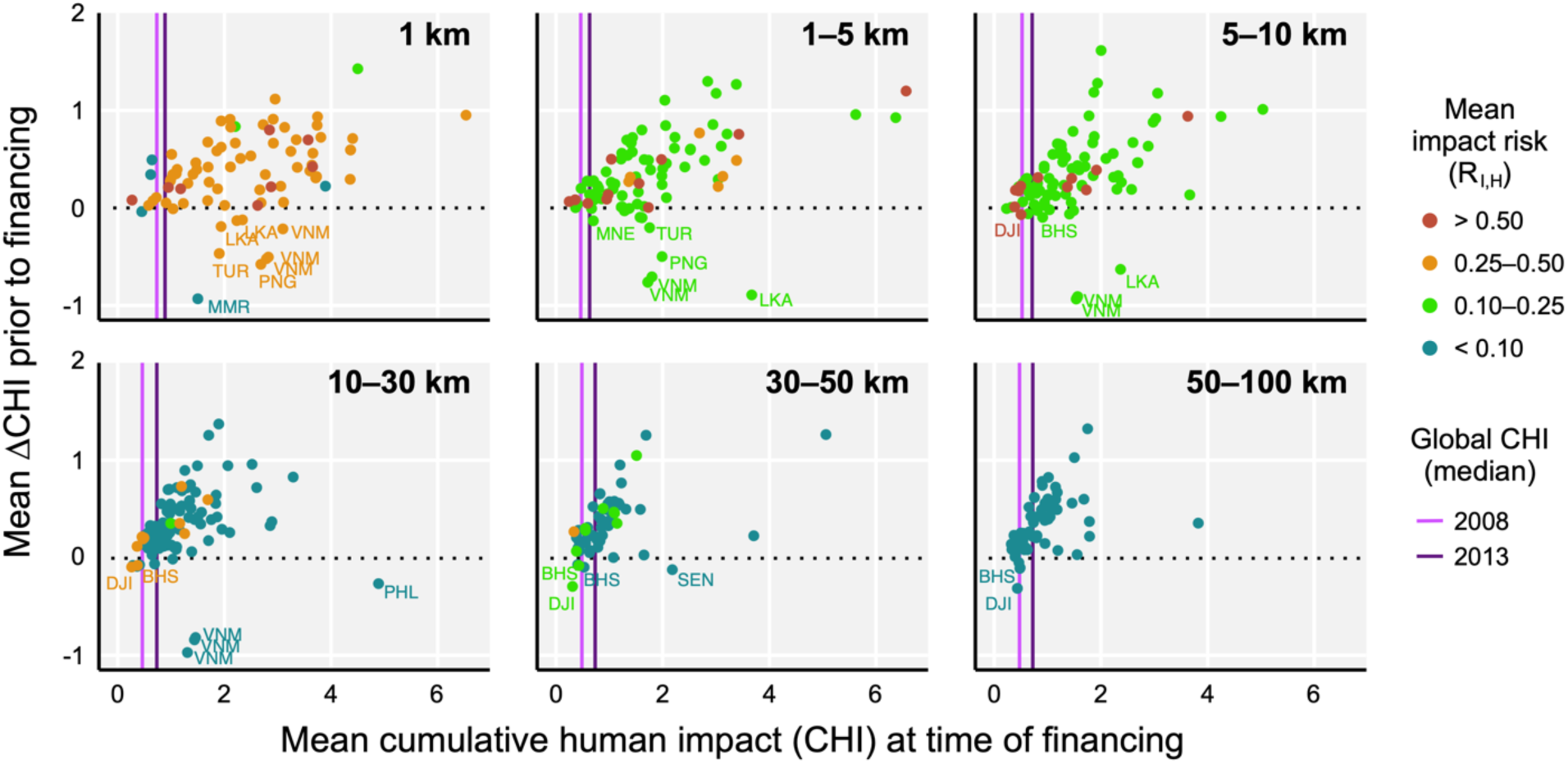
Mean impact risks of DFI projects at increasing distances relative to the state of marine ecosystems (cumulative human impact, CHI) and trends in their level of degradation (ΔCHI) prior to Chinese financing commitment. Projects below the dotted line (ΔCHI < 0) indicate prior reductions in CHI (i.e. ecosystem improvements) before financing. Vertical lines reflect the median CHI value in 2008 (light purple) and 2013 (dark purple) of all marine ecosystems within the respective distance range from global coastlines. Select projects of interest are labelled by country ISO code.

### Risks to sensitive socio-ecological features

As they do with habitats, ports present the greatest overall impact risks for the sensitive socio-ecological features included in this study (i.e. threatened species, marine protected areas, likely critical habitats, and Indigenous-use seas), generally followed by power plants and other facilities (Extended Data Fig. 3). While there is greater variation in the degree of risk imposed by DFI projects on these sensitive features than on habitats (particularly for more localized risks), the relationship between risks to habitats and all sensitive features is driven predominantly by the collinearity between the risks to habitats and threatened species (Extended Data Fig. 4). Some notable projects include Eritrea’s Hirgigo power plant upgrade, which presents high localized risks to likely critical habitats within potential Indigenous-use seas, as well as Tonga’s National Road Improvement Program, which presents high risks to nearby marine protected areas in addition to likely critical habitats and Indigenous-use seas. Overall, 27% of projects present high mean impact risks (*R_I_* > 0.25) to only one sensitive feature, while 25% and 4% present high impact risks to two or three types of sensitive features, respectively, most often within 1 km of the project site. See Supplementary Figs. 2-5 for regional maps of impact risks to all sensitive socio-ecological features.

All Chinese DFI projects in this study present some degree of impact risk to threatened species (Extended Data Fig. 1b). Ports present the greatest impact risks to threatened species (Fig. 4a), and impact risks decline substantially beyond 10 km from the project site. Within this 10 km distance, some variation is present between taxonomic groups. For example, the magnitude of risk posed by roads, bridges, and power plants tends to be greater for bony fishes, and the light pollution from transmission lines poses a far greater risk to sea birds than other taxa (Fig. 4b). Overall, the most at-risk taxa vary over distance. Sea birds are at greatest risk within 1 km of the project site, and bony fishes are the only taxonomic group where mean impact risks increase up to 10 km from the project site (Extended Data Fig. 5).

**Fig. 4.**
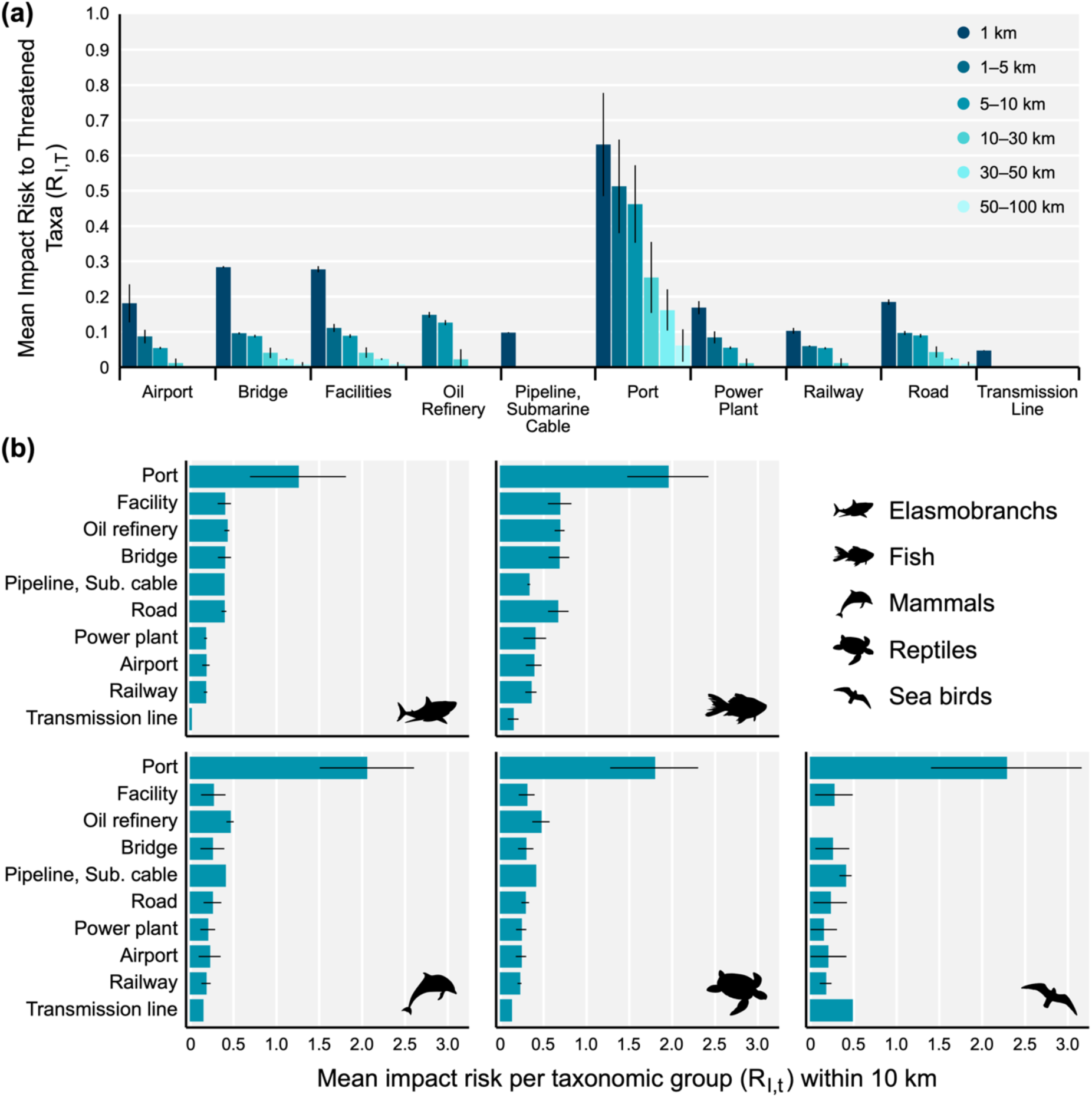
**(a)** Mean impact risk to all threatened taxa (*R_I,T_*) for different types of DFI projects at increasing distances from the project site. **(b)** Mean impact risk per taxonomic group (*R_I,t_*) for different types of projects within 10 km of the project site. Error bars represent the standard deviation of the mean.

Overall, 46% of DFI projects present exposure risk to MPAs, 31% to Indigenous-use seas, and 96% to likely critical habitats. In some cases, however, these risks may be difficult to avoid due to countries’ coastal characteristics that influence the likelihood that any coastal project may impact these sensitive areas. The majority of DFI projects in Cameroon, for example, present relatively high impact risks to Indigenous-use seas, but more than 75% of Cameroon’s coastline consists of important seas for coastal Indigenous communities (Fig. 5), so such high risks could be expected at any location along the coast. Other countries, however, deviate from what might be expected by chance. For instance, 89% of DFI projects in Vietnam and 55% of projects in Angola present moderately high impact risks to the few areas of likely critical habitat present within 10 km of their coastal waters (0.10 < *R_I,H_* < 0.25). In general, overlaps of project risks with these sensitive areas are close to what we would expect by chance alone for most countries; however, more countries start to present greater risks to these areas than we might expect when larger distances are considered (Fig. 5).

**Fig. 5.**
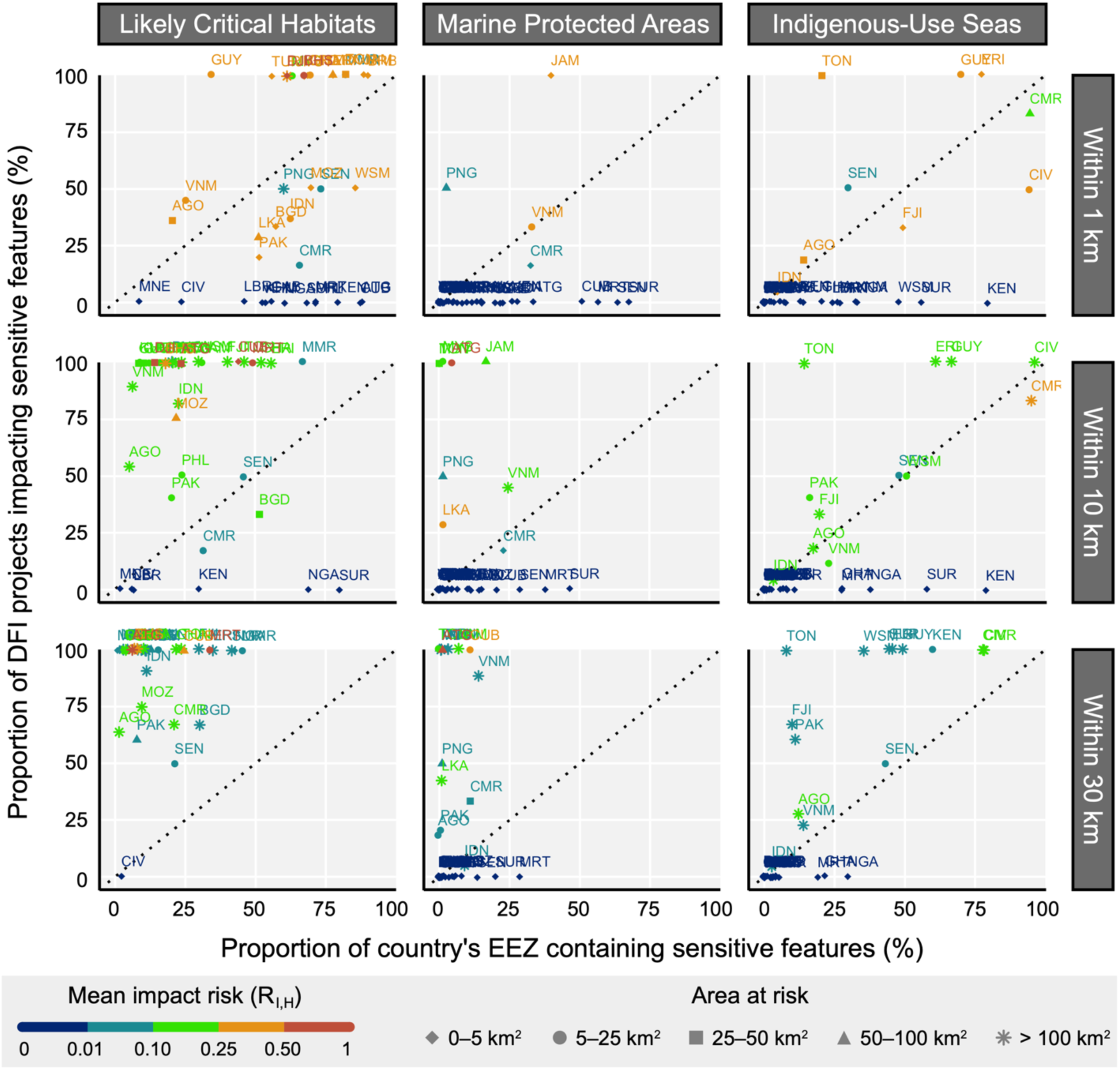
The proportion of each country’s DFI projects with impact risks to likely critical habitats, marine protected areas, and indigenous-use seas within 1, 10, and 30 km of the project site compared to the proportion of the country’s Exclusive Economic Zone (EEZ) containing the sensitive features within 1, 10, and 30 km of the coast. Countries above and below the dotted line indicate a higher and lower (respectively) proportion of projects with impact risks to sensitive features than would be expected at random. Countries are labelled by ISO code, with colour denoting the mean impact risk to habitats (*R_I,H_*) of all their DFI projects, and shape denoting the total ocean area under the mean impact risk. A positional jitter of 0.01 has been applied to enhance visibility of overlapping points.

While many projects present some exposure risks to these sensitive areas, the potential impact risks remain relatively small for most of these projects (Extended Data Fig. 6). Of those projects with exposure risks to marine protected areas, Indigenous-use seas, and likely critical habitats, only 6%, 9%, and 11%, respectively, present high mean impact risks (*R_I,H_* > 0.25) beyond 1 km from the project site. Some projects present high impact risks even up to 30 km from the site, such as the Hambantota Deep Sea Port in Sri Lanka (*R_I,MPA_* > 0.63), St. John Port in Antigua and Barbuda (*R_I,LCH_* > 0.49), and the Kribi Port Project in Cameroon (*R_I,IUS_* > 0.31). In total, 16,330 km^2^ of marine protected areas, 87,042 km^2^ of Indigenous-use seas, and 95,170 km^2^ of likely critical habitats face non-negligible impact risks (*R_I,H_* > 0) from DFI projects—of which 216 km^2^ (1.32%), 2739 km^2^ (3.15%), and 2648 km^2^ (2.78%), respectively, face high impact risks (*R_I,H_* > 0.25)— predominantly in Western Africa, Oceania, and the Caribbean (Extended Data Fig. 7).

Impact risks are present within potentially important seas for 34 groups of Indigenous peoples across 55 coastal communities, predominantly in Africa (65%), followed by Asia (16%), Oceania (11%), and South America (7%) (Fig. 6a). On average, 63% (1530 km^2^) of these communities’ Indigenous-use seas are at risk from DFI projects, with communities in Western and Central Africa facing the greatest mean impact risks, such as those in Côte d’Ivoire, Cameroon, and Angola (Fig. 6b). The implications of these risks are even greater considering that coastal Indigenous communities in Western and Central Africa tend to have some of the highest estimated annual seafood consumption rates. Communities in Côte d’Ivoire, in particular, face some of the highest impact risks to their seas and have some of the highest rates of seafood consumption (Fig. 6b).

**Fig. 6.**
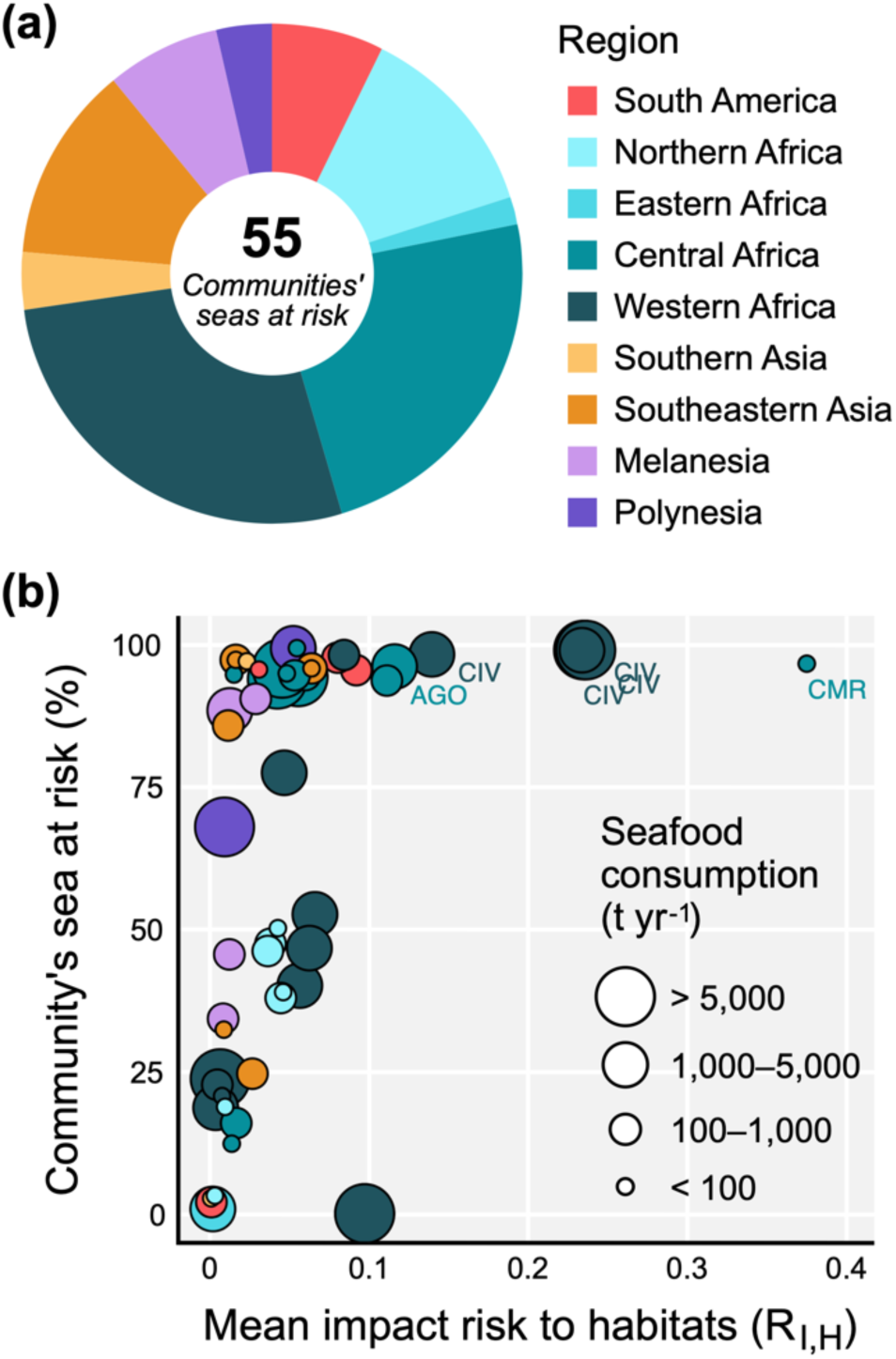
**(a)** Regional representation of the 55 coastal indigenous communities facing some impact risk from DFI projects. **(b)** Relationship between the proportion of each community’s indigenous-use sea (IUS) at risk and the mean impact risk to habitats (*R_I,H_*) within those at-risk seas. Colour indicates the geographic region, and point size indicates the estimated total annual seafood consumption of the community (in tons per year). Communities with the greatest impact risks are labelled by country ISO code.

## Discussion

### “Bluing” China’s overseas development finance

Our results echo previous concerns^25^ over the potential impacts of port developments financed by Chinese DFIs on marine socio-ecological systems, and these risks are likely to be even greater across the entire breadth of the BRI’s Maritime Silk Road. To date, considerations of the BRI’s impacts on marine systems have centered on these new ports and shipping routes, largely ignoring the potential indirect impacts of other land-based infrastructure financed by China. We found that coastal power plants, roads, bridges, and other facilities also present high risks to nearshore marine habitats from the effects of pollution from surface run-off, plastics, light, and noise. Unlike ports, these types of projects present more localized risks that diminish rapidly beyond 5–10 km from the development site, which may make it easier to manage or mitigate potential impacts over these smaller marine areas. While the critical habitats included in this study have yet to be verified on the ground, it is concerning that nearly all projects present some level of impact risk to these areas that are likely to meet the International Finance Corporation’s Performance Standard 6 criteria, which is intended to protect habitats of exceptional ecological value^28^. However, it is encouraging that few projects pose high risks to nearby marine protected areas, which are highly susceptible to the effects of human activities outside of their management boundaries^29^, though concerns remain for some marine protected areas in Antigua and Barbuda, Cameroon, and Sri Lanka (Extended Data Fig. 7).

An agenda is needed for “bluing” the BRI and all of China’s overseas development finance. Chinese banks have been criticized for not implementing the same level of biodiversity safeguards in their lending as other institutions, such as the World Bank^30^, which may be an important reason why their overseas development finance portfolio tends to place greater risks on terrestrial biodiversity and Indigenous lands than similar projects financed by the World Bank^22^. Although China’s policy banks require that host countries provide pre-loan environmental impact assessments for review, there is no standardized approach for vetting these assessments against international best-practices^30^. In practice, this can lead to rushed and insufficient impact assessments in countries with poor environmental governance, further undermining the CDB and CHEXIM guidelines that do exist^31^. Moreover, the limited accessibility of these assessments to the public contradicts the principles of the Aarhus Convention, to which several BRI countries are Party^32^, and limits transparency of the anticipated risks associated with coastal development projects.

China is well-positioned to extend its domestic environmental risk management policies to its overseas lending practices. China’s ecological conservation redline (ECRL) policy^33^, for example, has provided an important, data-driven framework for implementing ecosystem-based land use planning. The BRI’s multilateral platform presents an optimal opportunity for China to extend its expertise in institutional capacity building developed from the design of its ECRL policy to BRI member countries through collaboration, technical assistance, and codevelopment of harmonization programs. This strategy would help achieve China’s vision of building a national and global ecological civilization^34^, which is hindered by China’s current approach of deferring to host countries’ environmental standards for DFI project management. Extending China’s domestic experience to their finance partners would allow their DFI projects to meet the same (or greater) standards as their domestic environmental policies, which could help reduce the potential impacts of the projects it finances overseas. “Bluing” China’s overseas development finance will require greater integrated land-sea risk mitigation and management approaches^35^. Host countries should justify proposed coastal development projects within a mitigation hierarchy framework^36^ and provide a transparent, comprehensive, data-driven, and participatory-based assessment of land- and marine-based impacts. Chinese DFIs should further apply domestic and/or international standards to identify moderate- or high-risk projects, and negotiate with host countries to co-develop a long-term management plan to avoid, mitigate, remediate, or offset projected impacts to people and nature. At the very least, safeguards should set restrictions not just on projects directly overlapping with sensitive socio-ecological areas, but also on projects within close proximity to these features, such as marine protected areas, existing or potential marine critical habitats, and coastal Indigenous communities, to reduce the indirect impacts of land-based infrastructure on important marine systems.

The 114 Chinese DFI projects assessed in this study (those within 1 km of the coastline or a river within 100 km of the coastline) constitute one-fourth of the projects financed by the China Development Bank and Export-Import Bank of China between 2008-2019 that have been geolocated with high spatial precision^14^. However, it is likely that more coastal development projects have been financed that have yet to be mapped, as there are an additional 209 (31%) Chinese DFI loans where the location(s) of the projects have a spatial uncertainty of 25 km or more. The majority of China’s overseas development finance portfolio may thus have negligible potential risks to marine systems, or at least lower risks than those considered in this study. Additionally, several of the ports, airports, power plants, and other facilities financed by Chinese policy banks constitute expansions or upgrades of existing infrastructure, and our results show that the majority of DFI projects pose additional risks to marine ecosystems that are already facing relatively high human impacts. It is likely that the impacts of these projects will be smaller in magnitude than projects built in more intact ecosystems, though there may still be an ecological tipping point that has yet to be reached in heavily developed areas that these new projects could encroach upon^7^. For example, the Abidjan Port expansion in Côte d’Ivoire and ML-1 Railway expansion in Pakistan introduce additional risks to marine habitats that are some of the most degraded of all habitats included in this study; while these risks may not pose a threat to pristine ecosystems, researchers should be concerned if increased development could spur an ecosystem collapse. More research is needed locally to investigate the magnitude of risk posed by these development projects in heavily degraded habitats.

Estimating land-based risks to marine ecosystems remains a significant challenge for environmental impact assessments, as there are no standard models for capturing the effects of land use change, extreme weather, socio-economic feedbacks, and other uncertainties that are crucial for informing integrated land-sea management actions^35^. This reduces the consistency with which the direct and indirect risks of coastal infrastructure can be estimated between proposals, as well as the effectiveness of impact assessments to adequately capture the breadth of risks from the complex land-sea dynamics in coastal communities. Our global risk assessment is not a replacement for local risk assessments at each project site, which can more appropriately assess the potential impacts of development using more precise parameters to characterize the climatic, hydraulic, socio-economic, and operational features that ultimately influence the spread and magnitude of risk to marine socio-ecological systems. Additionally, the scarcity of temporal maps of marine habitats increases the difficulty of measuring trends in marine habitat loss. We used the best available maps from 2008^27^ as a historical reference for the earliest DFI projects in our database, but local risk assessments will be needed that can more precisely map the current/historical extent of marine habitats around development sites. Chinese DFIs can utilize a multitude of resources for proactive planning of their coastal and marine development projects; new groups are emerging to guide investors, policy-makers, and institutions on the environmental risks of infrastructure investments, such as the Taskforce on Nature-Related Financial Disclosures (TNFD), and international advisory boards are providing sustainable development recommendations to China, including the China Council for International Cooperation on Environment and Development (CCICED).

Socio-economic safeguards must also play a central role in “bluing” Chinese overseas development projects. We identified 55 coastal Indigenous communities, most notably in Western Africa, whose nearby coastal waters are at risk from the negative impacts of development, and this is likely a conservative estimate of the livelihoods that may be jeopardized from the loss of several marine ecosystem services^37^, such as the provision of seafood or the income generated from artisanal fisheries and tourism. For example, several Indigenous communities surrounding development projects in Côte d’Ivoire are estimated to consume more than 1000 tons of seafood per year, yet the habitats they rely on are facing some of the greatest ecological risks from development. China’s overseas development projects have a history of social conflict and protest^38^, and conflicts are already commonplace in coastal seas between competing users and resource uses^39^. Marine policy has struggled to fully incorporate the social sciences into evidence-based decision making^40^, and as more countries increase political recognition of coastal Indigenous communities’ rights over their historical marine resources^41^, there is an urgent need to ensure Chinese DFIs consider the implications of overseas coastal development through a lens of social equity and responsibility. Proactive planning should be inclusive of the voices of local communities and Indigenous peoples, and social impact assessments should be carried out alongside environmental impact assessments to ensure development does not diminish the rights, agency, or well-being of local and Indigenous communities.

### Directions for research and policy

This study is the first to incorporate global habitat and species vulnerability estimates to stressors associated with different types of coastal development projects in China’s overseas development finance portfolio. Our spatially explicit approach, which accounts for differences between habitats, taxonomic groups, and different types of development activities, can be used to estimate similar impact risks across other development finance portfolios, such as those from the World Bank, which are largely absent in the global coastal development literature. However, more local and regional risk assessments are necessary for estimating risks with greater precision than is possible at a global scale, and for those projects that have already been completed, post hoc impact evaluations should be implemented to track the realized effects (both positive and negative) of these development activities. Our results emphasize the urgent need for more in-depth investigations into Chinese DFI projects in Africa and the Caribbean, where port developments present exceptionally high risks, and especially in Western Africa, where risks are greatest for coastal Indigenous communities’ marine resources. Future investigations should be interdisciplinary, addressing not just the ecological challenges of coastal development in these regions, but also the social, cultural, economic, and political challenges in these unique contexts. Finally, studies are emerging that quantify the risks from new shipping routes and increasing traffic along the Maritime Silk Road on economic growth^42^, maritime accidents^23^, and carbon emissions^43^, yet the risks to vulnerable habitats and species have yet to be quantified and analyzed in depth. In addition to the features considered in this study, more research is needed investigating how the Maritime Silk Road may impact species’ behaviors, migration patterns, and the spread of invasive species.

President Xi Jinping has repeatedly stressed the importance of a green BRI, and China’s development of its ecological redline policy illustrates their capacity to deliver on national commitments for improving environmental protection and marine spatial planning^44^. Such national commitments and emerging screening tools should be extended to their overseas development finance portfolio. For example, the Foreign Economic Cooperation Office of China’s Ministry of Ecology and Environment has developed an Environmental Risk Screening Tool (ERST) for lenders and investors to consider location-based risks for potential projects, including local biodiversity^45^. The risk factors shown in the ERST process can be incorporated into broader project risk assessment through processes such as the “Traffic Light System,” developed in collaboration with the BRI International Green Development Coalition^45^, which would attribute risk categories to projects based on expected impacts to local biodiversity and emissions. With these new tools, Chinese DFIs can continue to support crucial infrastructure projects abroad while mitigating risks to local marine biodiversity. Importantly, China is hosting the final meeting of the 15^th^ Conference of the Parties to the Convention on Biological Diversity in April 2022, and there is an increasing global recognition of the importance of green investments, nature’s valuation in financial disclosures, and the integration of biodiversity values into development processes. China’s overseas lending should be guided by international commitments to sustainable development, ensuring that these activities can provide benefits to all people. However, China is just one of the key players in global development. Ultimately, social and environmental sustainability must be a cornerstone of all overseas development finance policy, whether by international organizations, such as the World Bank, or other countries, such as the “Blue Dot Network”^46^ launched by the United States, Australia, and Japan.

## Methods

### Data

#### Coastal and marine DFI projects

We obtained geolocated data for projects financed by China’s development finance institutions (DFIs)—the China Development Bank (CDB) and Export-Import Bank of China (CHEXIM)—from 2008 to 2019 from the Chinese Overseas Development Finance (CODF) database^47^. The CODF dataset contains 669 loans for projects represented as points (e.g. single structures), lines (e.g. roads, cables), or polygons (e.g. structural complexes, reservoirs, extraction areas) of varying levels of spatial precision. We included projects where the exact location is known (precision = 1), as the remaining projects have a location uncertainty of 25 km or greater^14^. For each project included in the analysis, we expanded upon the existing sector classification to distinguish and group the most similar types of projects according to their potential impacts on marine systems. Projects with the explicit goal of environmental protection, such as the Palisadoes repair project in Jamaica, were excluded from the analysis.

To identify DFI projects that may directly or indirectly impact marine systems, we used the most recent, high-resolution data of global shorelines^48^ and river networks^49^ to select (1) all marine-based projects and (2) all terrestrial-based projects within 1 km (Euclidean distance) of coastlines and/or rivers around the world. For this analysis, we excluded any riverside projects where the shortest river path distance from the impacted riverbank to the coastline exceeded 100 km; because our estimates of potential marine impacts are limited to the spatial extent of existing vulnerability data (see further), these riverside projects must be within 100 km of the coastline as determined by the extent of the cumulative human impact dataset’s ocean mask layer^2^, which does not always align with the extent mapped by Sayre et al.^48^, particularly for large river deltas. However, the consequences of this misalignment in coastline extent on our impact results are negligible, as this only affects projects located farthest from the coast. When more than one loan was provided for a single project, we only included the earliest (or highest) loan to avoid duplicating risk estimates. This resulted in the exclusion of 5 duplicate loans—Project IDs 214, 247, 276, 690, and 945^14^. After removing duplicates and any project whose maximum dispersal of risk did not reach the ocean (see section *Calculating exposure risk*), a total of 114 unique projects were included in the final analysis, including 10 shipping ports, 1 fishing port, 6 bridges, 33 roads, 6 railways, 5 airports, 1 oil refinery, 36 power plants (coal, oil, gas, nuclear), 2 transmission lines, 1 pipeline, 2 submarine cables, and 11 other facilities (e.g. housing developments, stadiums, office buildings) (Fig. 1a, Supplementary Table 1).

#### Marine habitats and existing human impacts

We obtained maps of the distribution of 20 habitats used by Halpern et al.^27^ to quantify cumulative human impacts on the world’s oceans, including intertidal, coastal, and offshore ecosystems. These maps include the presence/absence of each habitat within approx. 1 km^2^ ocean cells, allowing multiple habitats to occupy a single cell. Although some marine habitat maps have recently been updated given improvements in remote sensing technology, we use the original habitat maps generated by Halpern et al.^27^ in 2008 to align with the earliest year for which we have data on China’s DFI projects. We also obtained maps of the existing cumulative human impacts (CHI) to each ocean cell from Halpern et al.^1^, which provides annual CHI maps from 2003 to 2013 for select stressors (see section *Impacts prior to financing*).

#### Threatened marine species

We used the latest data from Butt et al.^50^ on marine species’ vulnerability to anthropogenic stressors to identify risks to marine species (see section *Calculating impact risks*). We considered the following five marine taxonomic groups whose status and distribution have been most comprehensively assessed and for which mean vulnerability scores have been estimated: elasmobranchs, bony fishes, marine mammals, marine reptiles, and sea birds (see Supplementary Table 2 for taxonomic information). We obtained species range maps from the IUCN Red List^51^ for all species within the taxonomic groups. We limited inclusion to extant, native or reintroduced species classified as ‘vulnerable,’ ‘endangered,’ or ‘critically endangered’ (see Supplementary Table 3 for all search criteria). In total, the ranges of 126 species of elasmobranchs, 101 species of fish, 19 species of marine mammals, 14 species of marine reptiles, and 64 species of sea birds were within the maximum range of exposure risk for the 114 DFI projects and were included in the analysis (see Supplementary Table 4 for all species included).

#### Marine protected areas

We obtained spatial data on marine protected areas (MPA) from the April 2021 version of the World Database on Protected Areas^52^. We included all legally designated, adopted, or inscribed regional, national, or international protected areas classified as marine or coastal (i.e., terrestrial and marine) protected areas. We dissolved overlapping polygon boundaries and excluded any MPAs where boundaries have not been mapped (i.e. point data). Boundaries of other effective area-based conservation measures were not present in the dataset.

#### Likely critical habitats

We used spatial data of the global critical habitat screening criteria from UNEP-WCMC^53^. This map identifies terrestrial and marine habitats that are “likely” or have “potential” to qualify as critical habitat based upon their on-ground presence and alignment with one or more criteria outlined by the International Finance Corporation’s Performance Standard 6 (IFC PS6)^28^. While these habitats have yet to be confirmed on the ground, we include all marine areas identified as “likely” critical habitat (LCH) as a representation of potential globally significant ecological areas.

#### Indigenous-use seas

We obtained data on the locations of coastal Indigenous peoples from Cisneros-Montemayor et al.^26^, which included 1,924 points representing 611 unique communities around the world. Here, coastal Indigenous peoples include “recognized Indigenous groups, and unrecognized but self-identified ethnic minority groups, whose cultural heritage and socio-economic practices are connected to marine ecosystems that are central to their daily lives and key to their nature-culture dynamics and concepts of surroundings, language, and world views”^26^. We excluded all points where the exact location was uncertain (approx. 48% of all points). Because the exact boundaries of coastal Indigenous territories surrounding these points are unavailable, and due to frequent disagreements between this dataset and other existing polygonal maps of Indigenous peoples’ lands^54^, we do not know the true distribution of the community in the landscape, nor how much of the coastline might be used by each community. As an approximation, we followed the assumption of Halpern et al.^27^ that 25 km represents the largest distance most small-scale fishers might travel to reach the shoreline. Thus, we created a 25 km buffer around each point and considered all coastlines within the buffer zone as potentially important to the local community. In some instances, points were not located within 25 km of the coastline; for these communities, we selected a section of the coastline in which the nearest river flowed to.

For each community’s coastline of importance, we created a 25 km buffer—13.5 nautical miles (nm)—as an estimation of the ocean area that is most important to Indigenous fisheries and marine resources, which we refer to as ‘Indigenous-use seas’ (IUS) (Extended Data Fig. 8). This distance aligns closely with countries’ territorial sea zone (22.2 km; 12 nm) and is expected to characterize most types of small-scale fisheries at a global scale^55^, though some have been known to travel distances of 75 km or more depending on their equipment^56^. We do not imply that these seas are owned and/or managed by the local Indigenous community, nor should it be interpreted that the absence of coastal Indigenous communities or IUS boundaries imply certainty in the presence of Indigenous communities or the importance of the sea for Indigenous communities. This data reflects assumptions based on our best available knowledge at a global scale.

#### Seafood consumption

The dataset of coastal Indigenous communities^26^ includes estimated annual seafood consumption rates for each community. These estimates represent the direct consumption of fish, invertebrates, and other marine species, based on existing data (where available) or estimated with a meta-analytical value-transfer model (see Cisneros-Montemayor et al.^26^ for methodological details).

### Stressors associated with DFI projects

#### Habitat loss or degradation

Habitat loss and degradation from coastal and marine infrastructure is the most direct stressor for DFI projects. This stressor is most likely to result from the construction of bridges, land reclamation from some airports, installation of pipelines and submarine cables, and the construction and maintenance of ports (e.g. quays, berths, dykes, breakwaters, dredging). The risks of habitat loss associated with these projects, however, are highly localized to habitats in the immediate surroundings of the construction sites. While some ports can reach up to 5 km in size^25^, we assume that habitats within 1 km of the project site will be most at-risk from construction across the diversity of DFI projects, consistent with approaches in previous studies^2, 57^. In contrast, risks of habitat loss/degradation arising from the trawling practices of some demersal fishing vessels is more expansive. These vessels, which may increase in frequency surrounding new/expanding fishing ports, travel large distances, collecting fish and other species by scraping along the seabed and contributing to the degradation of benthic habitats^58^. Thus, in addition to the localized risks associated with ports, we assume that risks of habitat loss/degradation associated with the increase in destructive demersal fishing extends to benthic habitats 100 km from fishing ports, a distance consistent with previous risk assessments surrounding ports^25^.

#### Noise pollution

Anthropogenic noise affects the behavior and physiology of a variety of marine species, with acoustic-sensitive species particularly vulnerable to this stressor^59^. While noise pollution may not always be a long-term stressor (e.g. temporary noise during construction), several DFI projects may continue to increased noise pollution beyond the initial construction phase. Because sound travels farther in water, underwater noise pollution (e.g. noise from shipping vessels) presents a greater risk to marine species than noise above-water^60^, and this has been the primary source of marine noise pollution investigated in the literature^59^. For port projects, we assume noise pollution risk from cargo and fishing vessels may reach up to 100 km from the port for the most sensitive species, consistent with previous studies^25^. Impacts of above-water noise from other types of infrastructure, such as railways, roads, bridges, power plants, and other facilities, are under-investigated in the literature, with most studies measuring impacts in controlled, artificial environments^60^. Given the lower frequency of noise from these projects, we assume they present more localized risks to species, unlikely to expand beyond 1 km from the source. Noise pollution from airports, however, may reach farther distances as planes fly above the ocean. While comprehensive evidence is scarce, one study^61^ found engine noise intensity underwater was 20 times greater within 1 km of an airport than 10 km from an airport. Though impacts will be dependent on flight paths, ocean depth, and other ambient characteristics, we assume noise pollution risks are greatest within 5 km from these coastal airports.

#### Light pollution

Artificial light can have negative effects on species behavior, reproduction, and communication, with flow-on effects on communities and ecosystem functions^62^. However, few studies have measured impacts of light pollution on marine species over distance. Light pollution is typically measured according to light intensity, and different sources and types of light can lead to different intensities and subsequent impacts^63^. Given these complexities, we assume additive risks from light pollution scale with the size of infrastructure. For roads, bridges, and power transmission lines where streetlights or tower lights may be lower in intensity, we assume risks from light pollution are unlikely to extend beyond 1 km, as has been observed in some birds^64^. For power plants, airports, and other large facilities that demand more light per area, light pollution may extend farther from the site given the greater light intensity of the project and the potential spread of artificial skyglow^63^. In the absence of published studies measuring the distance at which these individual projects may impact species, we assume risks are greatest within 5 km of the site. Light from shipping/fishing vessels is also a potential threat for species further from the coast, which may be exacerbated through port developments. In line with previous approaches^25^, we assume light pollution surrounding ports may be significant up to 100 km from the port due to increased sea traffic at night.

#### Thermal pollution

Increases in water temperature (i.e. thermal pollution) downstream of power plants are a prevalent concern across the world^65, 66^. Temperature increases can reach up to 10 °C depending on the design of the plant and aquatic factors influencing the diffusion of the heat effluent^66, 67^. While the European Union’s standard thermal pollution limit is a 3 °C increase, acceptable limits may be as small as a 1.5 °C increase depending on the type of habitat^66^. Several studies have created their own models of thermal pollution dispersion from individual power plants^68–70^, but these require several parameters—such as local tide and current dynamics, as well as wind, water, and outflow velocities—for which global data (where available) are not of sufficient detail for modeling thermal plume dispersion from individual plants. However, several studies indicate that the most significantly impacted areas tend to occur within 5 km of the discharge site^69–72^. Thus, we considered thermal pollution risks from coal, gas, oil, and nuclear power plants to reach up to 5 km from the project site.

#### Inorganic pollution

Inorganic pollution is a significant concern for coastal ecosystems facing intensive urban development. These pollutants are often the result of urban runoff, depositing metals and minerals leached from impervious surfaces into nearby waterways, such as concrete, asphalt, and other building materials^73^. Risks from inorganic pollution are the most common across DFI projects that increase impervious surface area adjacent to coastal waters, including roads, bridges, railways, power plants, oil refineries, airports, ports, and other facilities. Hydrological transport models are often needed to robustly estimate the dispersal of these pollutants, as risks are influenced by numerous factors like water and wind velocity, tides and currents, depth, advection, and the partition coefficients of metals^74^. Compared to thermal pollution, the dispersal potential of chemical pollutants appears to be more variable, with some primarily concentrated within just a few kilometres of the outflow^75^ and others maintaining ecologically-significant concentrations more than 20 km from the coastline^76, 77^. To estimate generalized exposure risks of inorganic pollution, we defined a decay function based on the dispersal of pollutants in river plumes mapped by the ocean cumulative human impact dataset^27^, which assumes exposure risk declines non-linearly over a distance of 30 km (see section *Diffusive risks*).

#### Nutrient and organic pollution

Pollution from nutrient-rich compounds, such as those found in fertilizers, are a severe threat to aquatic ecosystems. These nutrients often enter rivers and coastal waters through agricultural runoff and sewage/wastewater from treatment plants or ships, which can lead to eutrophication, harmful algal blooms, or hypoxia in the most affected areas^78^. Other organic pollutants, such as those in herbicides/pesticides from agricultural runoff and petroleum from ships and oil refineries, also present considerable risk to aquatic ecosystems, as many of the associated hydrocarbons, organochlorines, and other compounds can be toxic to aquatic species^79, 80^. For the DFI projects included in this study, ports present the most significant risk of nutrient pollution, and both ports and oil refineries present the greatest risk of organic pollution. We use the same equation of diffusive risks from inorganic pollution to estimate exposure risks of nutrient and organic pollution surrounding these project locations.

#### Sedimentation

Sedimentation is a prominent threat for rivers and coastal ecosystems around the world, which can be exacerbated by land use changes for agriculture, mining, and urban development^81^. While most DFI projects considered in this study may increase the short-term risk of sedimentation during construction, ports present the most prominent long-term risk of sedimentation from regular dredging^82^. Like the organic and inorganic pollutants described previously, estimating the dispersion of sediment particles is complicated by local hydrological conditions, as well as additional factors unique to sedimentation, such as grain size, cohesiveness, and sinking/fall velocity^74^. While these additional factors ultimately require individual modelling of sedimentation dispersion potential, for this analysis, we assume sedimentation risk follows the same decay function as organic and inorganic pollutants.

#### Plastic pollution

Pollution from floating plastic debris (including microplastics) is one of the most pervasive stressors in the sea, capable of spreading thousands of kilometers from the initial source^83, 84^. Plastic litter is the most probable cause of plastic pollution associated with DFI projects, including litter from roads, bridges, ports, and other facilities^2, 50, 85^. Although plastic litter can travel across entire oceans, we limit the extent of this potential risk to 100 km surrounding these DFI projects to capture the pervasive nature of plastic pollution while limiting the range at which we expect plastic density to be greatest for a single project.

#### Invasive species

Invasive species may be introduced through a variety of mechanisms, but one of the most common for marine environments is through the transport of ballast water in shipping vessels^86^. A global standard for estimating invasion risk for different types or characteristics of vessels is still lacking, with some studies modeling the spread of invasive species up to 1000 km from the port based on the estimated volume of ballast water released at a given port^27^. However, because most invasive species thrive in shallower waters^2^, we assume exposure risks are greatest immediately surrounding shipping ports, where ballast water transfers will be concentrated, and become negligible for most invasive species beyond 100 km from the port.

#### Wildlife injury

Increased marine traffic carries additional risks to inadvertent injury of marine wildlife through vessel strikes^27^. This risk differs from other stressors that result in the intentional loss or injury of wildlife, such as biomass removal or dynamite/cyanide fishing (poisons/toxins). Some studies have estimated risks of vessel strikes within 50 km of ports^25^, while others expect risks to be present along global shipping routes^27^. Because the density of shipping lanes can still be high 100 km from a given port, we assume risks of wildlife injury will be non-negligible up to 100 km from ports.

#### Biomass removal, bycatch, entanglement, poisons and toxins

Fishing vessels are associated with several stressors that are unique from shipping vessels, including biomass removal, bycatch (biomass removal of non-target species), entanglement from nets and other fishing gear, and the use of poisons or toxins (e.g. dynamite or cyanide fishing)^50^. For most of these stressors, these risks are prevalent across most types of commercial fisheries, which may increase with the creation or expansion of fishing ports. Because commercial fishing vessels travel large distances, we assume the risks of biomass removal, bycatch, and entanglement are expansive, spanning up to 100 km from the port’s location, consistent with our other risk estimates from marine vessels. The use of poisons or toxins, however, are more common in artisanal fisheries. These small-scale fishers can often be classified into different typologies based on the distance from the shore that they travel^55^. Though there is variation between typologies, the majority of artisanal fishers travel less than 30 km from the coast. Thus, thus we assume that risks from all fishing-related stressors associated with artisanal fisheries are greatest within 30 km from fishing ports.

### Vulnerability to stressors

#### Threatened species

We use data from Butt et al.^50^, which estimates the average vulnerability of each of the five taxonomic groups included in this study to the following 15 stressors relevant for DFI projects: habitat loss/degradation, noise pollution, light pollution, thermal pollution, inorganic pollution, nutrient pollution, organic pollution, sedimentation, plastic pollution, invasive species, wildlife injury, biomass removal, bycatch, entanglement, and poisons/toxins (Supplementary Table 5). Vulnerability weights range from 0 to 1 and represent the expected average vulnerability for all species within the taxonomic group based upon the vulnerability of species’ biophysical, movement, reproduction, specialization, and spatial traits to different stressors estimated through extensive literature reviews, expert elicitation, and sensitivity tests. Although important variations still exist between species in the same taxonomic group, we use this global trait-based vulnerability framework to provide a *relative* index of vulnerability among diverse taxonomic groups to a variety of anthropogenic stressors at a global scale. See Butt et al.^50^ for methodological details.

#### Marine habitats

We use data from Halpern et al.^27^, which estimates the vulnerability of all 20 habitats included in this study to the following 15 stressors relevant for DFI projects: light pollution, inorganic pollution, nutrient pollution, ocean pollution, sea surface temperature, direct human pressure, benthic structures, shipping, invasive species, demersal destructive fishing, demersal non-destructive fishing (high and low bycatch), pelagic fishing (high and low bycatch), and artisanal fishing. For simplicity, we grouped all non-destructive fishing stressors together to reflect “commercial fishing” that is unique from demersal destructive fishing and artisanal fishing. Vulnerability weights for this commercial fishing stressor thus reflect the mean vulnerability weights of the four non-destructive fishing stressors. Like the taxa vulnerability index, habitat vulnerability scores reflect *relative* vulnerability among marine ecosystems to different stressors based upon the literature and expert elicitation. Original vulnerability weights could reach a potential maximum of 5, but we rescaled these weights to fit the same 0 to 1 range as the taxa vulnerability weights (Supplementary Table 6). See Halpern et al.^27^ for methodological details.

### Calculating exposure risks

Each type of DFI project presents different risks to marine habitats and taxa, including different stressors and scales of potential impact. Here, we define exposure risk as the likelihood that a given ocean cell may be exposed to a particular stressor with non-negligible impacts on the habitats or taxa occurring within the cell. For all DFI projects, we assume risks are greatest within 1 km of the project site and decline with distance. To capture differences in the distribution of risk of different projects, we generated several risk indices that approximate the dispersal of exposure risk from each site (see Extended Data Table 1 for a summary of exposure risk distances by DFI project for each of the 15 stressors). All distances were calculated in ArcGIS under the World Mollweide projection, using Euclidean distances and accounting for land barriers. See section *Stressors associated with DFI projects* for justification of the spread of exposure risks for different projects and stressors.

#### Localized risks

In several instances, DFI projects present highly localized risks to habitats and taxa within their immediate surroundings. All projects associated with the risk of *habitat loss or degradation*—bridges, pipelines, submarine cables, ports, and airports with seaplane terminals— present localized risks of this stressor within their construction zone (i.e. < 1 km, but see section *Expansive risks* for the extent of habitat loss risks from destructive demersal fisheries). Additionally, impacts of land-based *noise pollution* is unlikely to spread beyond 1 km of the source for most types of DFI projects, like railways, power plants, oil refineries, roads, bridges, and other facilities. Similarly, risks of *light pollution* from projects with a low light intensity—power transmission lines, roads, and bridges—are likely to be negligible beyond 1 km. For these project-stressor combinations where risks are highly localized, we attribute exposure risk as present (*R_E_* = 1) for ocean cells within 1 km of the project site, or absent (*R_E_* = 0) for all other cells.

#### Diffusive risks

Some projects, given their size or the nature of their stressors, present risks beyond their immediate surroundings. DFI projects contributing to more intense *light pollution*, including power plants, oil refineries, airports, and other facilities, may significantly affect aquatic communities up to 5 km from the project site. Given the high noise volume from airplanes, *noise pollution* from airports also has the potential to impact communities to a similar spatial extent. This distribution of risk is also consistent with existing evidence on *thermal pollution* of power plants. To characterize these diffusive risks, we estimated exposure risk for a given ocean cell as a function of distance (*x*) from the project site (in meters), where risk declines linearly up to 5 km from the site:

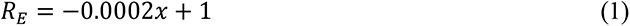

The distribution of risks from artisanal fisheries are unique from those of commercial fisheries. While commercial fishers often travel hundreds of kilometers from the coast (see section *Expansive risks*), this is not characteristic of small-scale fishers, often traveling less than 30 km from the coast. To estimate the risks posed specifically by artisanal fisheries at fishing ports (*biomass removal*, *bycatch*, *entanglement*, and *poisons/toxins*), we assume exposure risk declines linearly up to 30 km from the port:

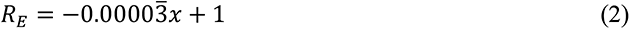

To estimate the distribution of risks from *sedimentation* and *inorganic*, *organic*, and *nutrient pollution*, we defined a logistic equation using parameter optimization^87^ to reflect the non-linear decline in exposure risk over distance based on maps of effluent dispersal in river plumes from the CHI dataset^27^:

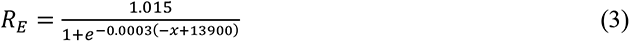

This function assumes pollution risks remain high within 5 km of the project site and decline rapidly beyond 10 km until approaching *R_E_* = 0 (Extended Data Fig. 9). At 30 km (*R_E_* = 0.008), additional reductions in risk become insignificant as the line approaches the asymptote. For simplicity, we assume pollution risks beyond 30 km from the project site are equivalent to zero.

#### Expansive risks

For some stressors, risks can span exceptionally large distances from the project site because of the nature of the stressor or the nature of the DFI project. *Plastic pollution*, for example, can spread thousands of kilometers with the ocean current, generating a significant risk to species and ecosystems far beyond the original source. Ports also increase the distribution of risks for stressors attributed to commercial shipping/fishing vessels, including *light* and *noise pollution*, *invasive species*, *wildlife strikes*, *biomass removal*, *bycatch*, and *entanglement*, as well as *habitat loss* specifically from destructive bottom-trawling practices of some demersal fisheries. For these stressor-project combinations, we assume the risk of exposure is greatest at the project site (i.e. where plastic debris first enters the ocean and where the concentration of shipping/fishing vessels is greatest) and decreases linearly up to 100 km from the site:

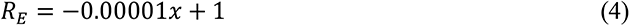

While this represents a conservative estimate of the true extent that plastic debris and commercial shipping/fishing vessels can travel (and thus impact marine systems), this range is consistent with other studies considering the distribution of these stressors in the ocean^2, 25, 85^.

### Calculating impact risks

#### Habitats

While exposure risk estimates the likelihood that a stressor may be present, some habitats will be more vulnerable to the effects of the same stressor exposure risk. Thus, we calculated the impact risk (*R_I_*) per habitat across all habitat stressors (*n* = 12) for each ocean cell as:

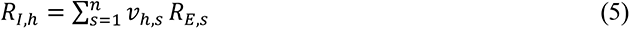

where *ν_h,s_* is the vulnerability weight of habitat *h* to stressor *s*, and *R_E,s_* is the exposure risk of stressor *s* for that cell, the product of which is the individual impact risk of stressor *s* on habitat *h*. Because each cell can contain multiple habitats, we focus our main results on the mean impact risk of all *n* habitats present in each cell:

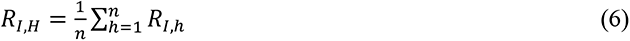

For easier comparability across impact risks, we rescaled all *R_I,H_* values within [0, 1] based upon the maximum observed impact risk across all cells (*R_I,Hmax_* = 2.321).

#### Sensitive socio-ecological features

To calculate impact risks for threatened taxa, we used the same approach as Eq. 5, replacing habitat vulnerability weights with taxon vulnerability weights (*ν_t,s_*) from Butt et al.^50^, across all species stressors (*n* = 15) and rescaling the mean impact risk of all taxa in each cell (*R_I,T_*) within [0, 1] based on the maximum observed impact risk (*R_I,Tmax_* = 4.000):

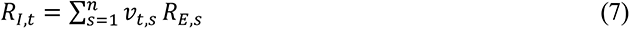

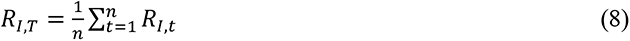

For sensitive territories—marine protected areas (MPA), likely critical habitats (LCH), and Indigenous-use seas (IUS)—we first created a binary indicator for the presence (1) or absence (0) of each territory in the ocean cell. When a sensitive territory is present, we consider the impact risks to be equivalent to the impact risks faced by the habitats within their boundaries (*R_I,H_*), such that:

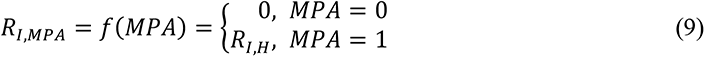

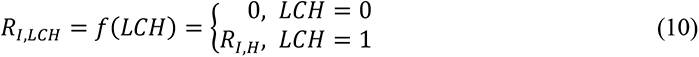

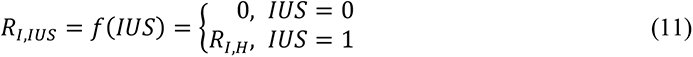

Finally, we generated a composite index reflecting the total impact risk for all sensitive features present in each cell (*R_I,F_*):

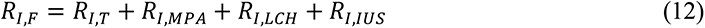

with a potential minimum *R_I,F_* = 0 (no sensitive features at risk) and maximum *R_I,F_* = 4 (all sensitive territories present, with the maximum potential impact risks on their habitats and threatened taxa).

#### Differences between habitats and taxa

While some of the habitat stressors are equivalent to the taxa stressors (e.g. light, inorganic, and nutrient pollution), several habitat stressors represent proxies of multiple taxa stressors. For example, the direct human stressor is a proxy for all stressors associated with coastal development and increasing coastal populations, including habitat loss/degradation, noise pollution, and plastic pollution, and the shipping stressor encapsulates the combined threats of noise pollution, plastic pollution, and wildlife injury/strikes^27^. Because the taxa-based stressors are more explicit and more numerous, we treat all habitat stressors as measures of the taxa stressors. For each habitat stressor, we determined if it had a similar singular equivalent taxa stressor, or if it encompassed multiple taxa stressors (Supplementary Table 7). Where the habitat stressor consisted of multiple taxa stressors, the exposure risk of the taxa stressors in that cell were reduced to reflect their contribution to the cumulative habitat stressor. For example, the impact risk to habitats from shipping was calculated as the product: 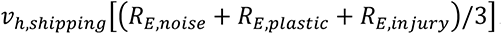. See Supplementary Table 7 for the treatment of each habitat stressor’s exposure risk relative to taxa stressors.

### Impacts prior to financing

#### Baseline ocean condition

To identify reference points of the level of ocean degradation prior to financing of each DFI project, we obtained annual stressor impact maps from the CHI dataset^1^ spanning 2008 to 2013. Annual impact data is only available for the following stressors included in our analysis: light pollution, nutrient pollution, organic pollution, shipping, direct human impact, sea surface temperature, demersal destructive fishing, demersal non-destructive fishing (high and low bycatch), pelagic fishing (high and low bycatch), and artisanal fishing. We added these stressor impact maps together to create our own CHI maps for each year between 2008 and 2013 that only include stressors associated with DFI projects, though this is a conservative estimate that does not include all land, ocean, and climate-based impacts on the ocean. We compare each DFI project with the CHI map corresponding to the year that the project was financed. Projects financed after 2013 are compared with the 2013 CHI map, which is the most recent data available. Because the CHI values are continuous, we also calculated the global median CHI for 2008 and 2013 at distances of 1, 5-10, 10-30, 30-50, and 50-100 km from the world’s coastlines to enhance interpretation of CHI values.

#### Trends in ocean condition

To determine the trajectory of change in CHI (ΔCHI) prior to financing of each DFI project, we subtracted the CHI for 2003^1^ from the CHI of years 2008-2013 (depending on year of financing). We followed the same methodology to generate the 2003 CHI map as the 2008-2013 CHI maps. For each ocean cell, ΔCHI > 0 indicates increasing human impacts prior to financing, and ΔCHI < 0 indicates decreasing impacts (i.e. improvements in ocean condition). Because there is variation in when projects were financed, ΔCHI values between projects are not always based on the same temporal range.

### Assumptions and limitations

While nearly all DFI projects present similar risks during construction—such as noise pollution, light pollution, organic/inorganic pollution, and sedimentation—these risks are often short-lived. Thus, we only consider long-term risks posed by stressors that have a lasting impact on marine environments, like continual pollution from run-off or the permanent loss of habitat in the foreseeable future. Additionally, we do not consider risks from stressors that are largely unknown or expected to be rare or unpredictable for a particular type of project, such as the risks posed by electromagnetic fields or thermal pollution from submarine cables^57^. We also do not account for interactions between stressors that may amplify risks of individual stressors, such as plastic pollution increasing the dispersal potential of invasive species^88^, nor do we account for the individual or additive impacts of climate change on DFI impact risks or the condition of the ocean prior to financing (e.g., ocean acidification, sea level rise, ultraviolet radiation). Risks will likely change when considering these additional factors that contribute to habitat quality, and future assessments should take into account more detailed information regarding the current condition of the ocean. Finally, we recognize that, in some industries, greater advances have been made to reduce the environmental footprint of development. We do not consider the mechanistic or operational characteristics of each DFI project, which may significantly improve their environmental performance (and thus reduce risk) beyond previous industry trends. More localized risk assessments must be conducted for individual projects that can account for these project-specific attributes, as well as the local demographic, socioeconomic, topographic, hydrological, and climatic characteristics that ultimately determine socio-ecological risks posed by coastal development.

## Acknowledgements

We acknowledge the Arapaho, Cheyenne, Ute, Massachusett, Chumash, Turrbal, and Jagera peoples, Traditional Owners of the land on which we live and work around the world, and pay our respects to their Elders past, present, and emerging. We are grateful to C. Klein, B. Halpern, and M. Frazier for supporting our application of their ocean and species datasets, as well as A. Cisneros-Montemayor and Y. Ota for providing us with data on coastal Indigenous communities. K.P.G. received funding for this project from the Climate and Land Use Alliance, David and Lucile Packard Foundation, and Rockefeller Brothers Fund.

---

**Extended Data Fig. 1.**
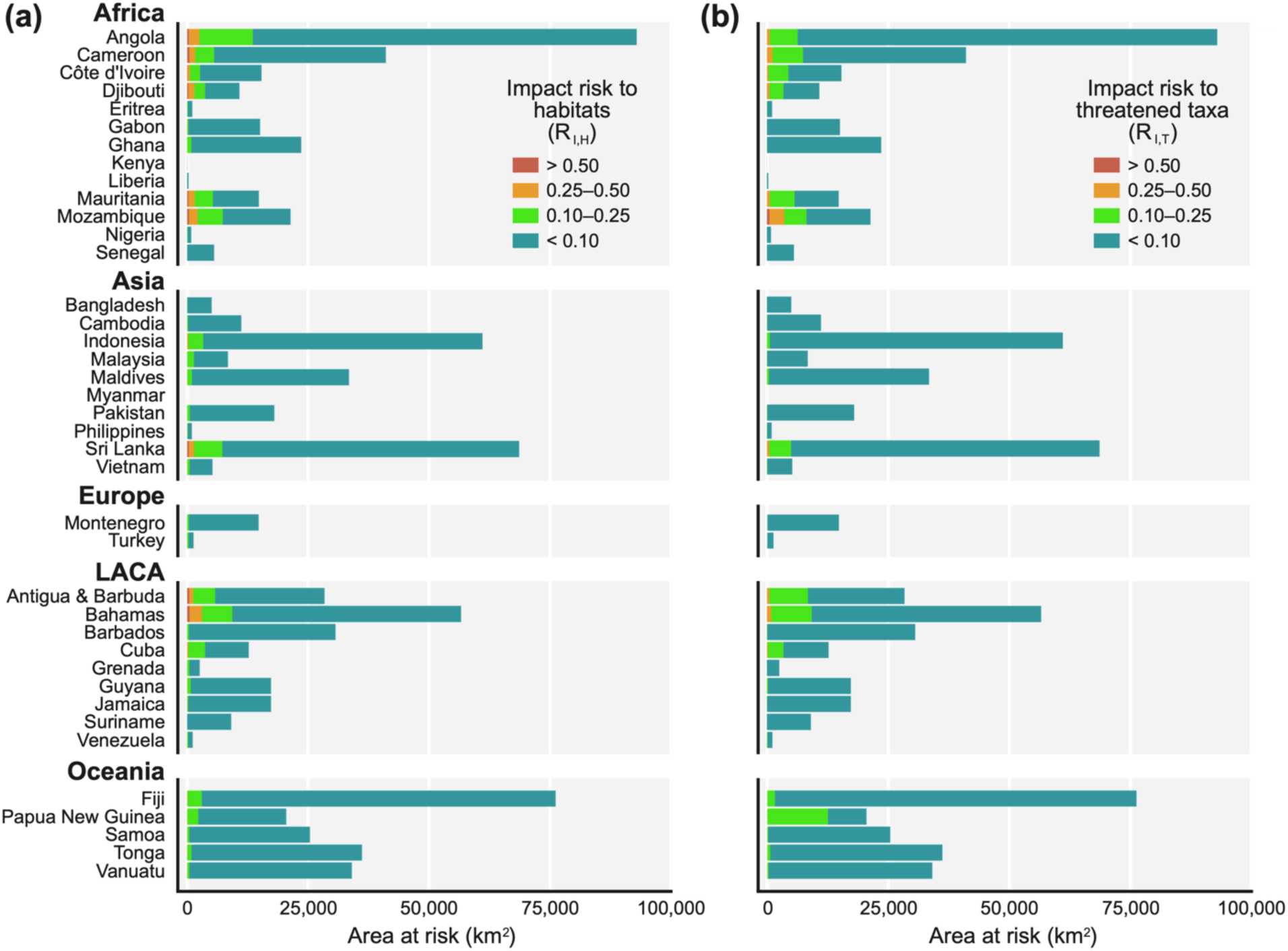
Total ocean area facing high and low impact risks to **(a)** habitats (*R_I,H_*) and **(b)** threatened taxa (*R_I,T_*) for the 39 countries with coastal development projects financed by Chinese DFIs. LACA = Latin America and the Caribbean.

**Extended Data Fig. 2.**
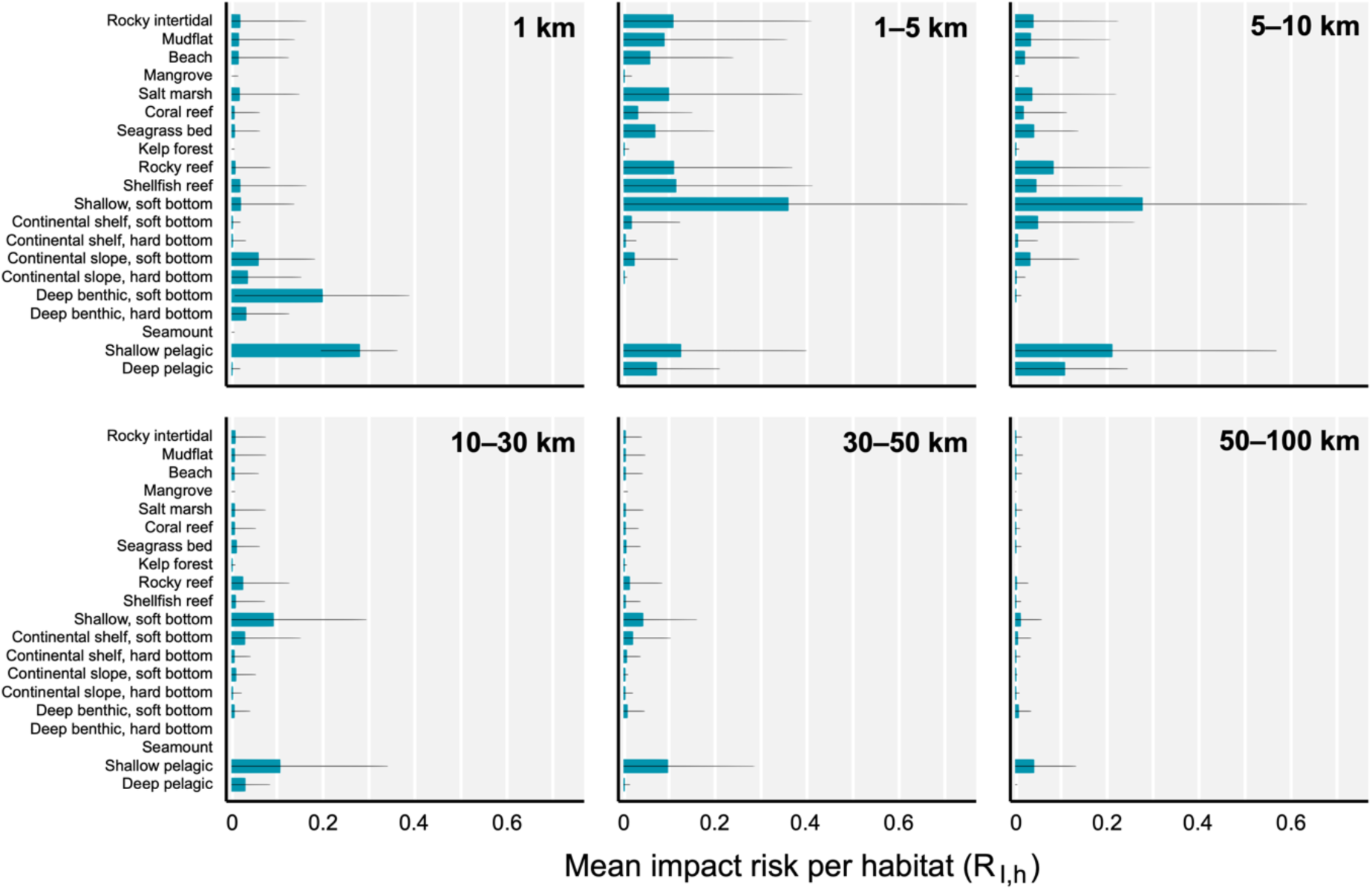
Mean impact risk to each habitat (*R_I,h_*) for all DFI projects at increasing distances from the project site. Error bars represent the standard deviation of the mean.

**Extended Data Fig. 3.**
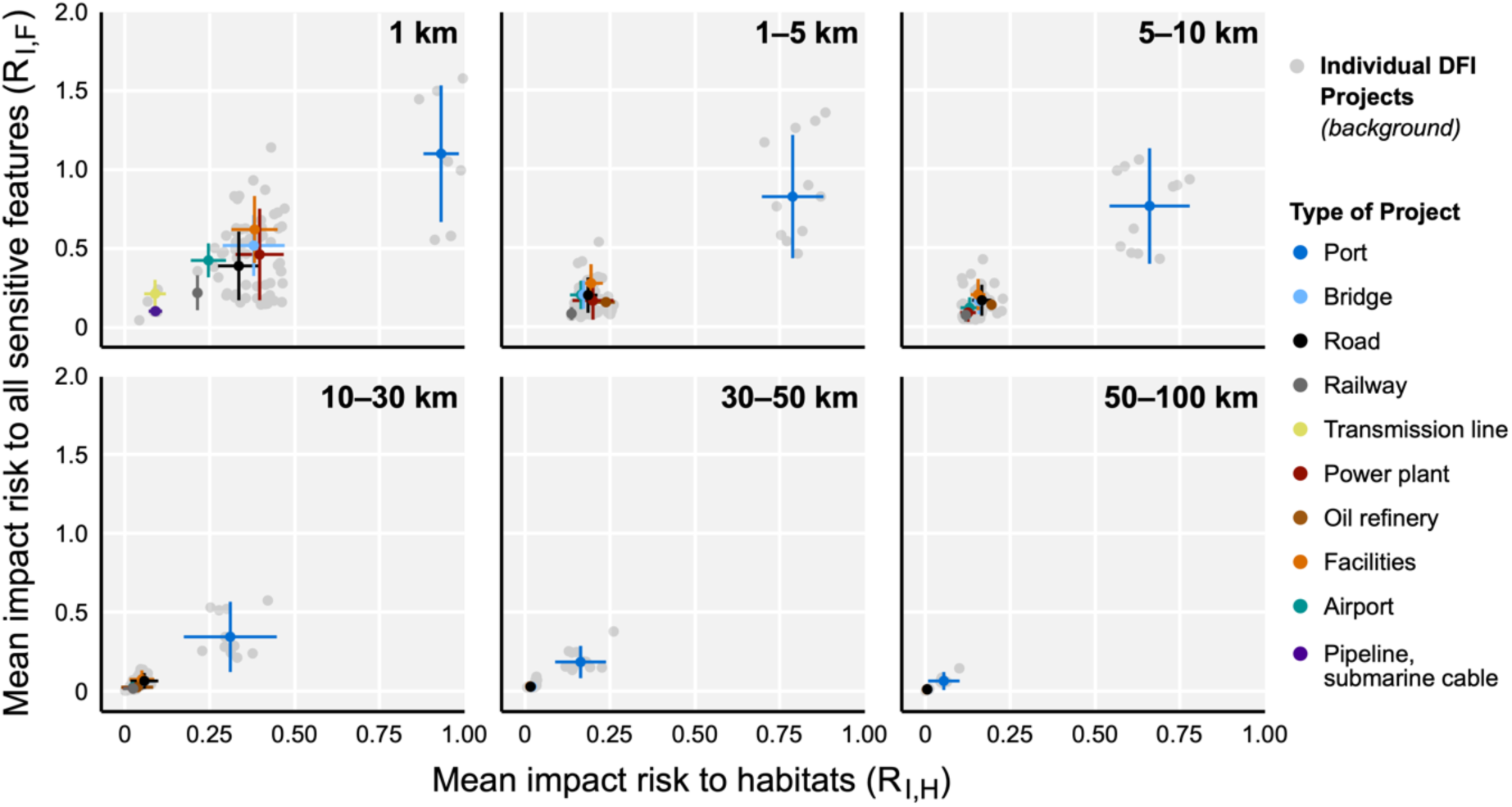
Relationship between mean impact risks to habitats (*R_I,H_*) and all sensitive socio-ecological features (*R_I,F_*) for different types of DFI projects at increasing distances from the project site. Error bars represent the standard deviation of mean impact risks for habitats (horizontal) and sensitive features (vertical) for each type of project.

**Extended Data Fig. 4.**
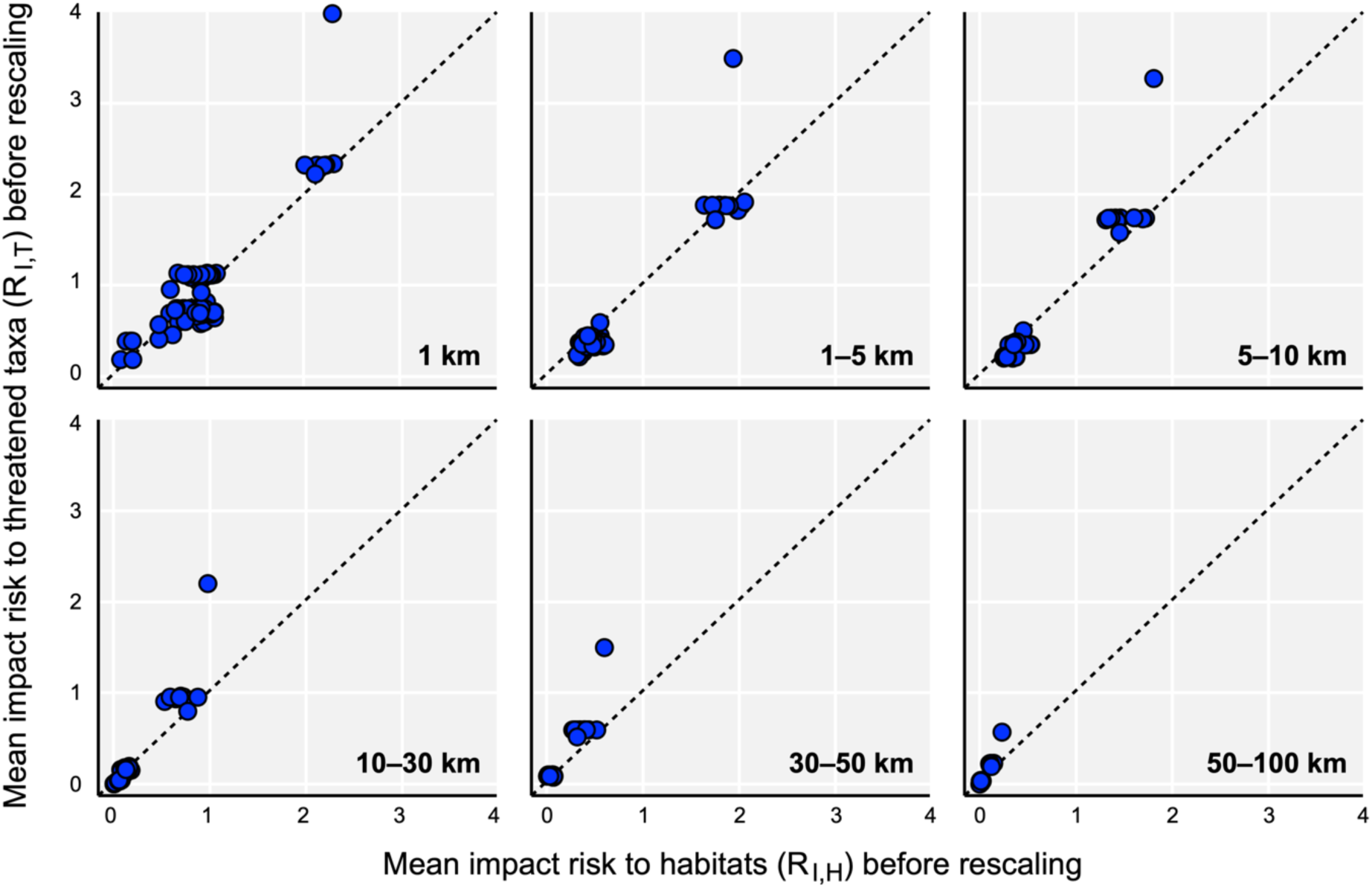
Relationship between mean impact risks to habitats (*R_I,H_*) and threatened taxa (*R_I,T_*) prior to rescaling each index for DFI projects at increasing distances from the project site. Dotted line shows a one-to-one relationship for reference.

**Extended Data Fig. 5.**
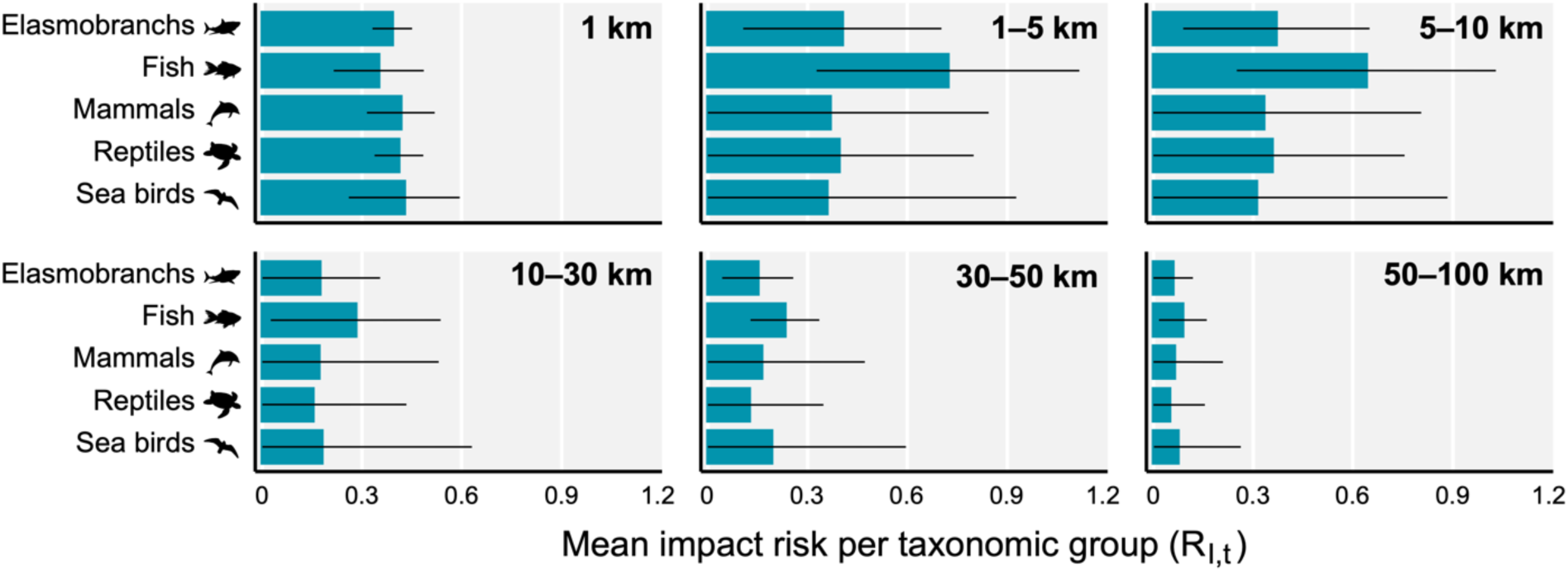
Mean impact risk to each threatened taxonomic group (*R_I,t_*) for all DFI projects at increasing distances from the project site. Error bars represent the standard deviation of the mean.

**Extended Data Fig. 6.**
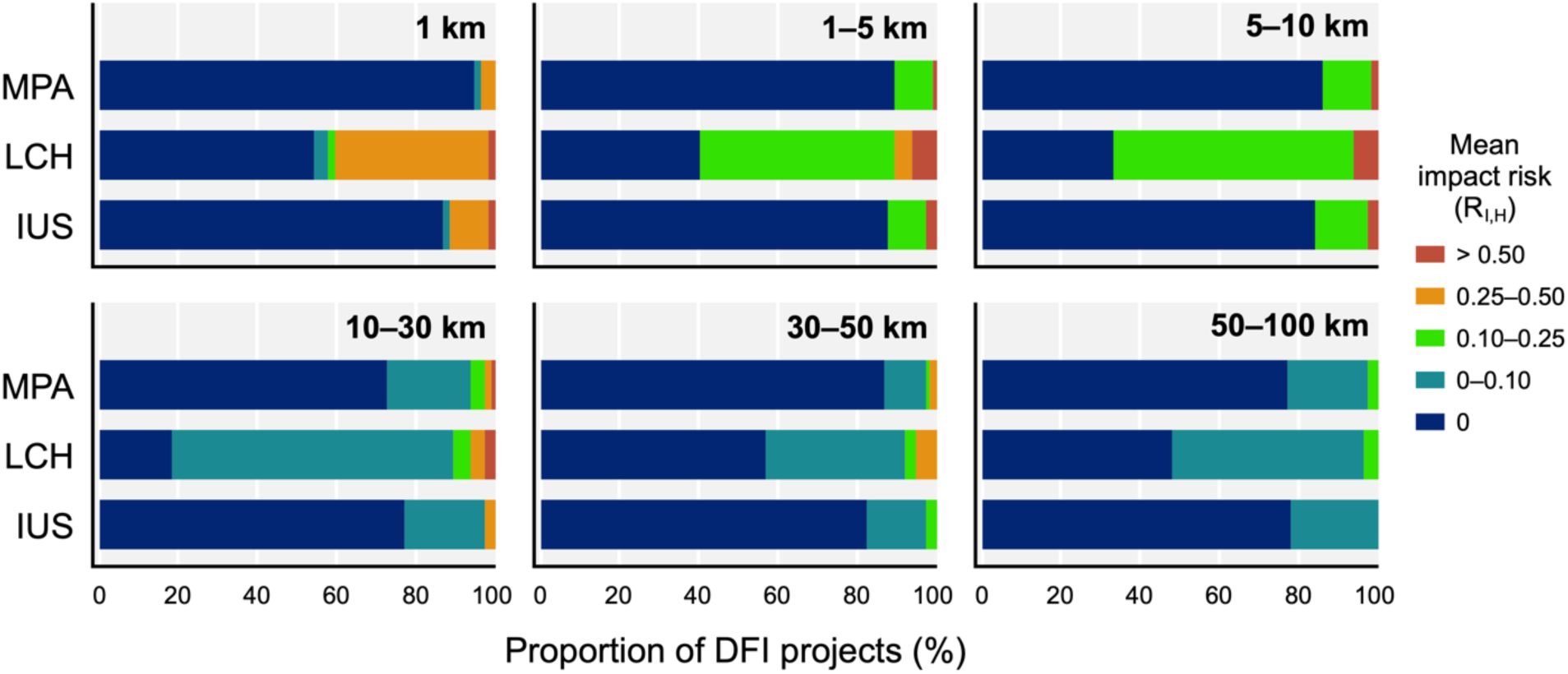
Proportion of all DFI projects presenting varying levels of impact risk to habitats (*R_I,H_*) within marine protected areas (MPA), likely critical habitats (LCH), and indigenous-use seas (IUS) at increasing distances from the project site.

**Extended Data Fig. 7.**
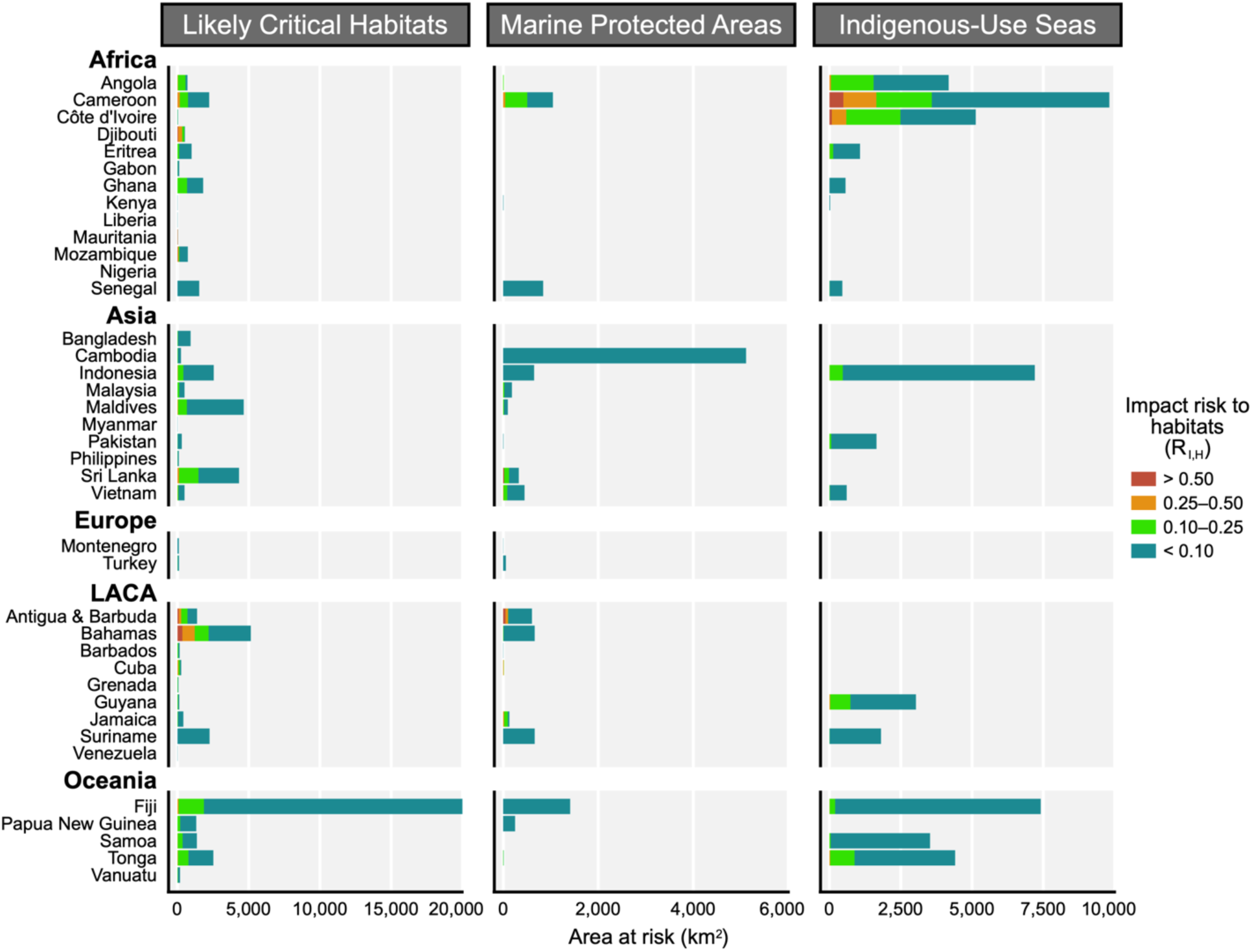
Total ocean area facing high and low impact risks to habitats (*R_I,H_*) within likely critical habitats, marine protected areas, and Indigenous-use seas for the 39 countries with coastal development projects financed by Chinese DFIs. LACA = Latin America and the Caribbean.

**Extended Data Fig. 8.**
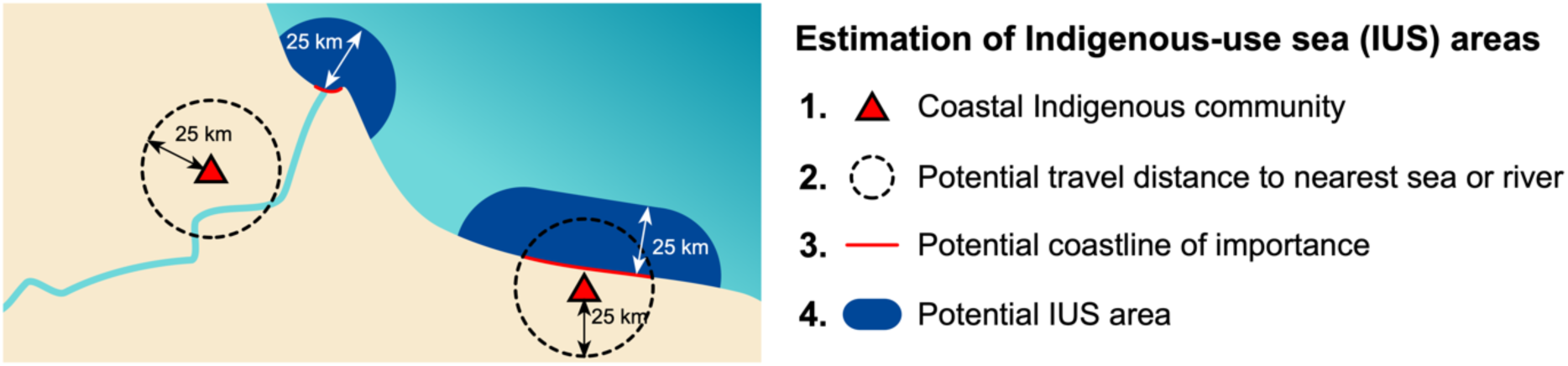
Illustrative example of the methodology for estimating Indigenous-use sea (IUS) areas based on point locations of coastal Indigenous communities. Map is a hypothetical illustration only.

**Extended Data Fig. 9.**
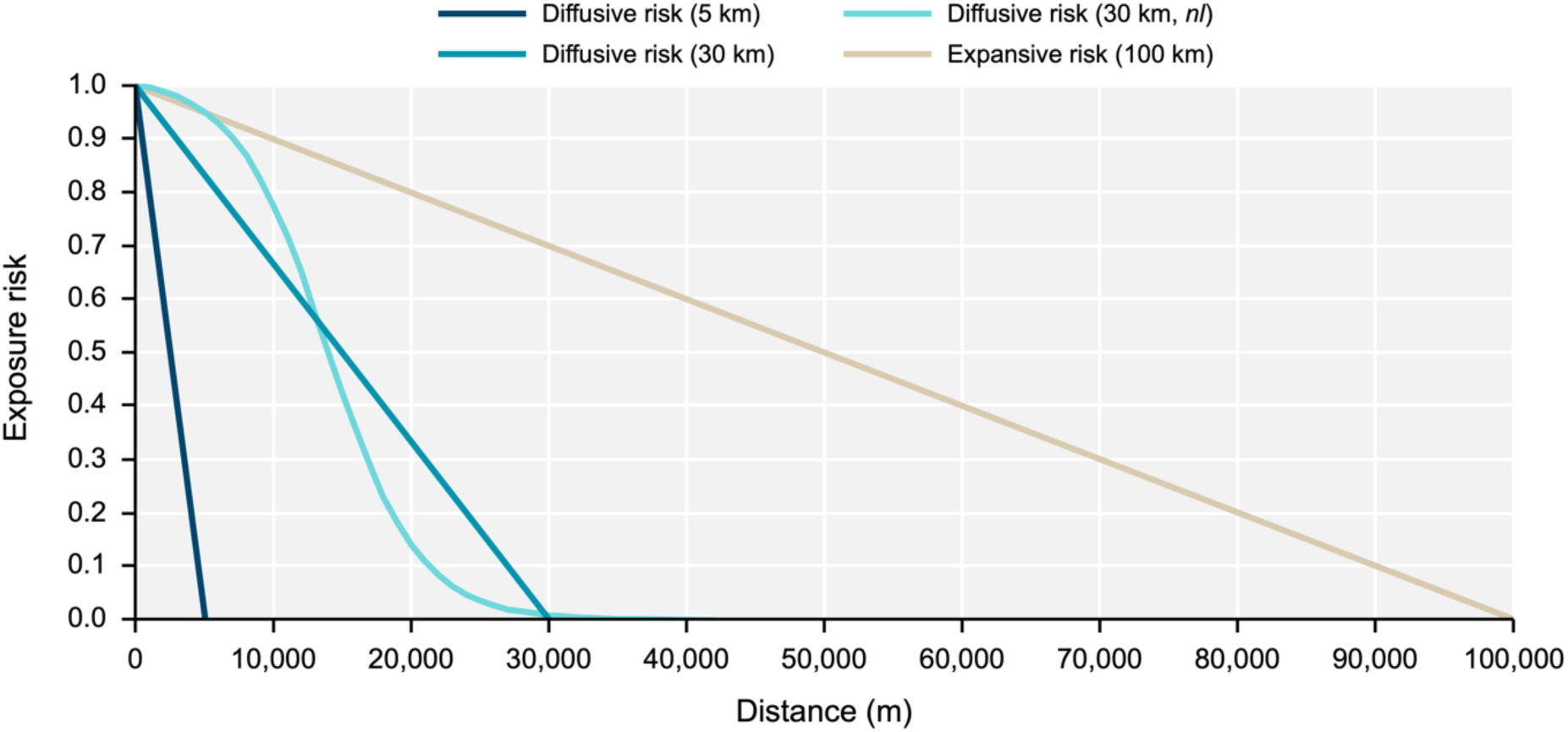
Functions defined to estimate the linear and non-linear (*nl*) decay of exposure risks (*R_E_*) as distance from the DFI project site increases. See Extended Data Table 1 for the stressor-project combinations for which each function is applied.

**Extended Data Table 1.** Stressors associated with each type of Chinese DFI project and the estimated distance of exposure risk from the project location. Localized risks (1 km) are represented as the presence (R = 1) or absence (R = 0) of risk. Risks over larger scales exhibit a linear decline in risk as distance increases, except for those marked as non-linear (*nl*). See footnotes for differences in risk distributions for different activities.

| Project type | Habitat loss | Noise pollution | Light pollution | Thermal pollution | Inorganic pollution | Nutrient pollution | Organic pollution | Sediment. | Plastic pollution | Invasive species | Wildlife injury | Biomass removal | Bycatch | Entangle. | Poisons, toxins |
| --- | --- | --- | --- | --- | --- | --- | --- | --- | --- | --- | --- | --- | --- | --- | --- |
| Transmission line |  |  | 1 km |  |  |  |  |  |  |  |  |  |  |  |  |
| Power plant (coal) |  | 1 km | 5 km | 5 km | 30 km (*nl*) |  |  |  |  |  |  |  |  |  |  |
| Power plant (gas) |  | 1 km | 5 km | 5 km | 30 km (*nl*) |  |  |  |  |  |  |  |  |  |  |
| Power plant (nuclear) |  | 1 km | 5 km | 5 km | 30 km (*nl*) |  |  |  |  |  |  |  |  |  |  |
| Power plant (oil) |  | 1 km | 5 km | 5 km | 30 km (*nl*) |  |  |  |  |  |  |  |  |  |  |
| Oil refinery |  | 1 km | 5 km |  | 30 km (*nl*) |  | 30 km (*nl*) |  |  |  |  |  |  |  |  |
| Railway |  | 1 km |  |  | 30 km (*nl*) |  |  |  |  |  |  |  |  |  |  |
| Road |  | 1 km | 1 km |  | 30 km (*nl*) |  |  |  | 100 km |  |  |  |  |  |  |
| Bridge | 1 km | 1 km | 1 km |  | 30 km (*nl*) |  |  |  | 100 km |  |  |  |  |  |  |
| Other facility |  | 1 km | 5 km |  | 30 km (*nl*) |  |  |  | 100 km |  |  |  |  |  |  |
| Airport | 1 km<sup>a</sup> | 5 km | 5 km |  | 30 km (*nl*) |  |  |  |  |  |  |  |  |  |  |
| Pipeline | 1 km |  |  |  |  |  |  |  |  |  |  |  |  |  |  |
| Submarine cable | 1 km |  |  |  |  |  |  |  |  |  |  |  |  |  |  |
| Port | 1 km<br>100 km<sup>b,c</sup> | 100 km | 100 km |  | 30 km (*nl*) | 30 km (*nl*) | 30 km (*nl*) | 30 km (*nl*) | 100 km | 100 km | 100 km | 100 km<sup>b</sup><br>30 km<sup>b,d</sup> | 100 km<sup>b</sup><br>30 km<sup>b,d</sup> | 100 km<sup>b</sup><br>30 km<sup>b,d</sup> | 30 km<sup>b,d</sup> |

<sup>a</sup> Specific to airports requiring land reclamation
<sup>b</sup> Specific to fishing ports
<sup>c</sup> Specific to destructive demersal fisheries
<sup>d</sup> Specific to artisanal fisheries

