## Supplementary Material for "Mapping the risks of China’s global coastal development to marine socio-ecological systems"

### Supplementary Figures

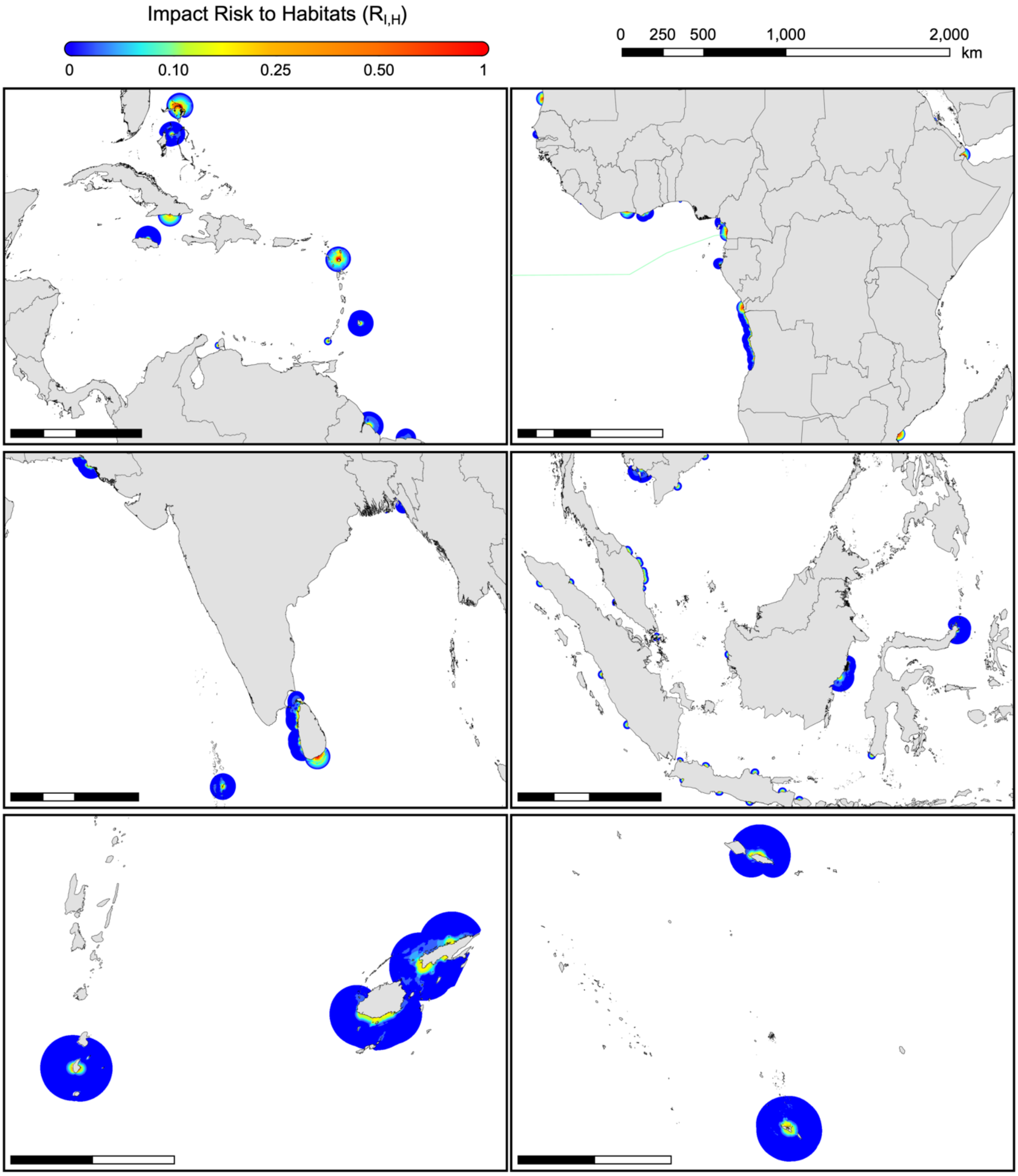


**Supplementary Fig. 1.** Larger regional maps of the distribution of impact risks to near- and off-shore habitats ($R_{I,H}$) from each DFI project site (see Fig. 1b).

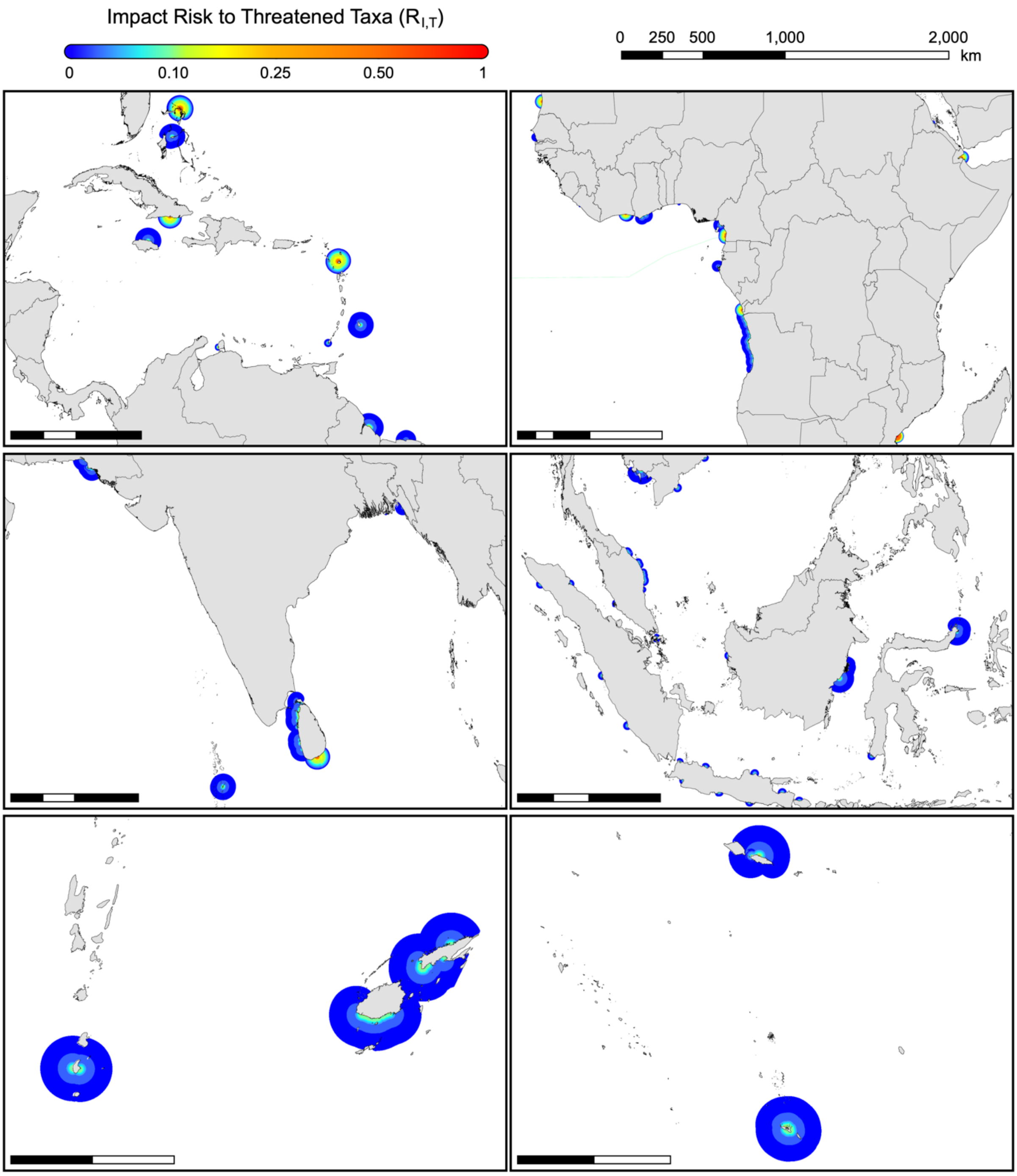


**Supplementary Fig. 2.** Regional maps of the distribution of impact risks to threatened taxa ($R_{I,T}$).

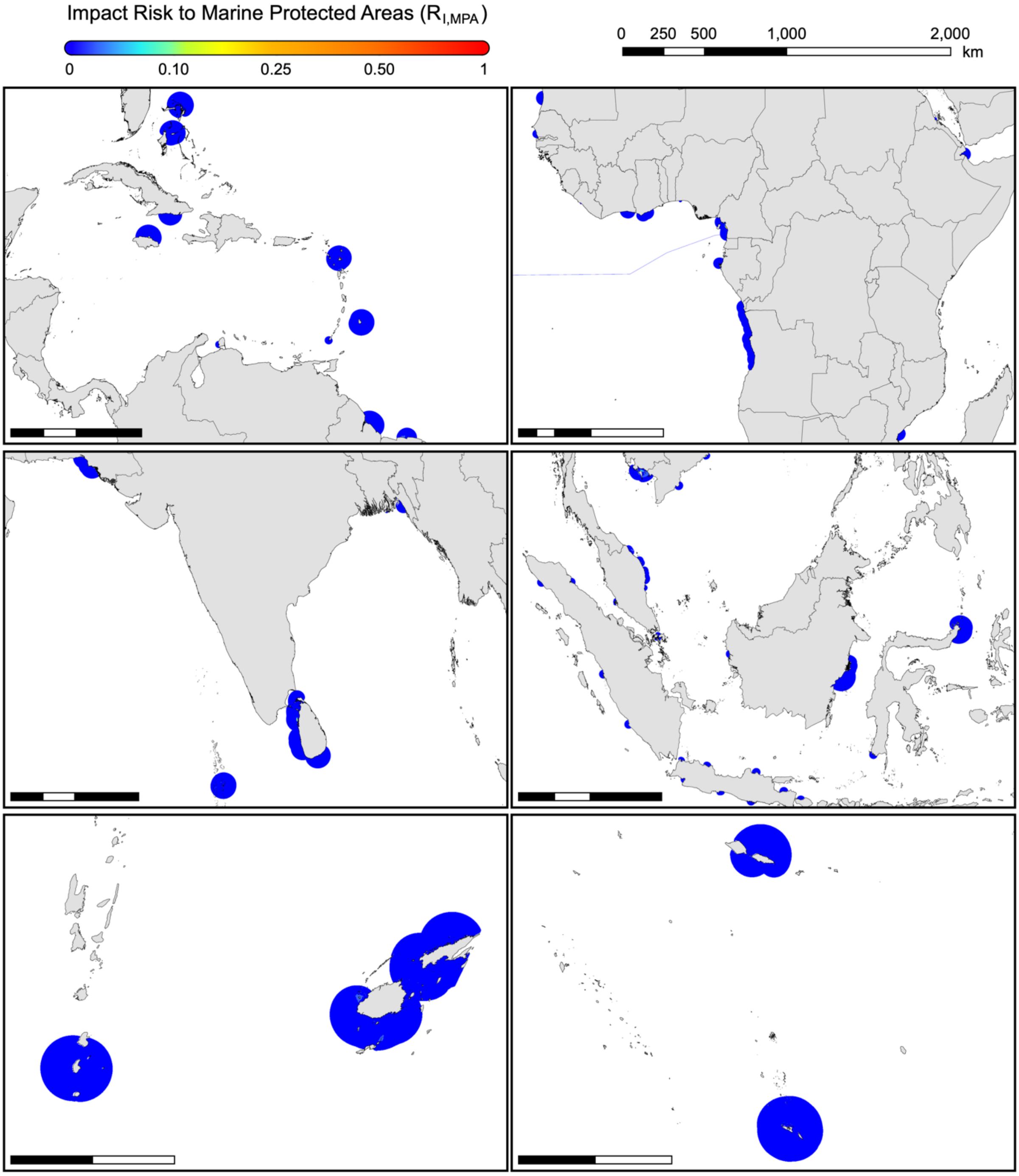


**Supplementary Fig. 3.** Regional maps of the distribution of impact risks to marine protected areas ($R_{I,MPA}$).

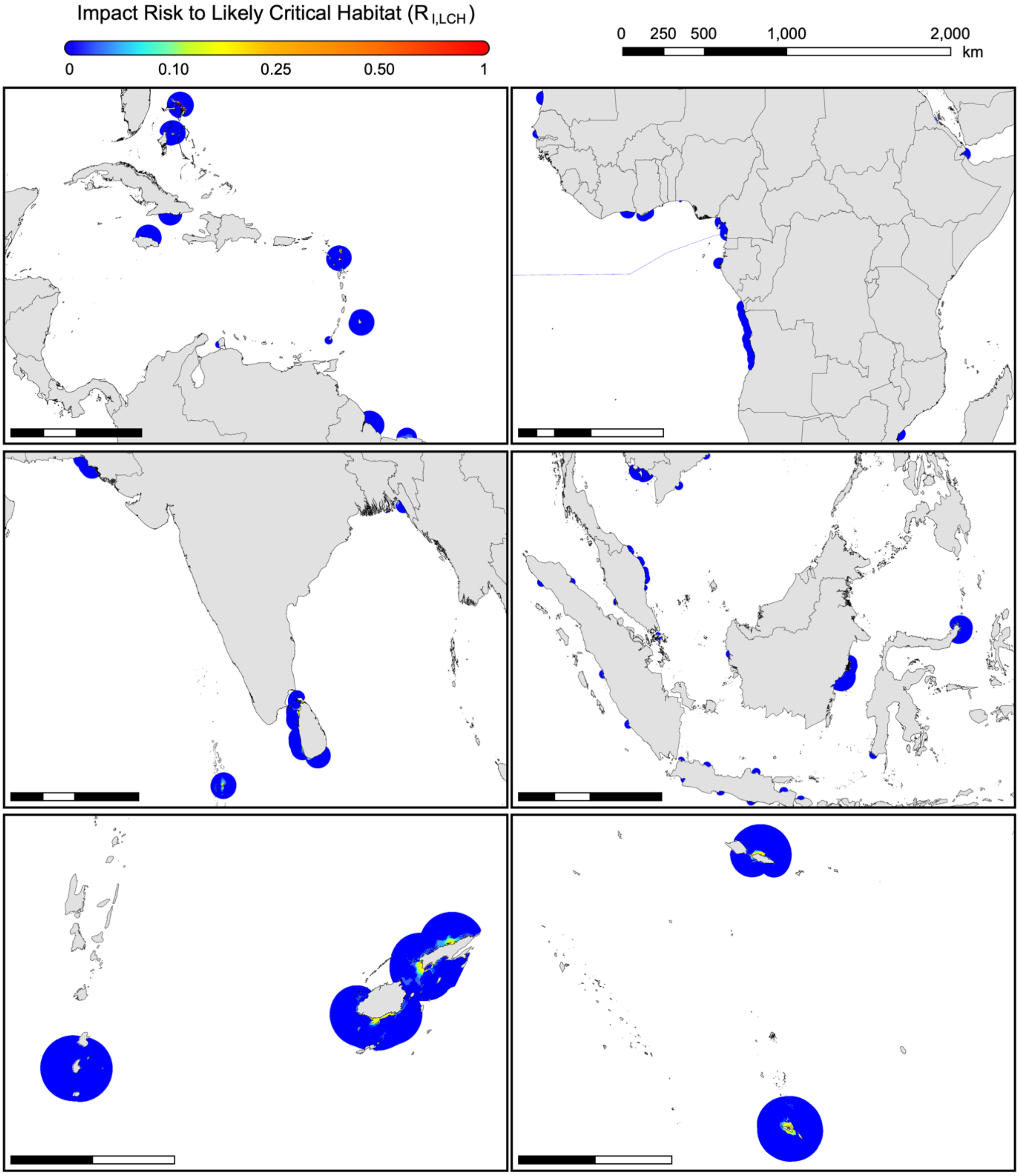


**Supplementary Fig. 4.** Regional maps of the distribution of impact risks to likely critical habitats ($R_{I,LCH}$).

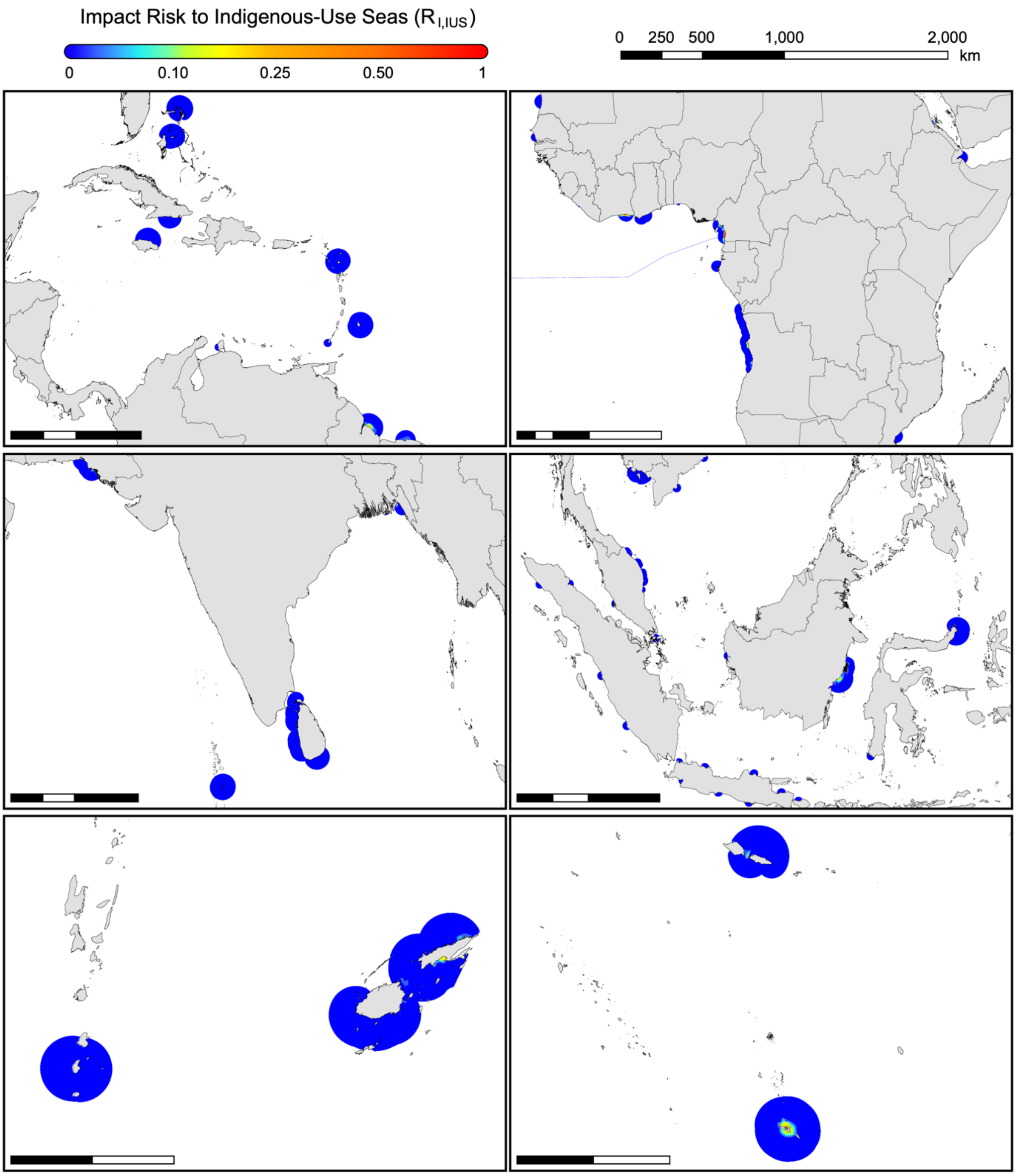


**Supplementary Fig. 5.** Regional maps of the distribution of impact risks to Indigenous-use seas ($R_{I,IUS}$).

### Supplementary Tables

**Supplementary Table 1**. Description of 114 projects financed by Chinese development finance institutions (DFIs) included in this study. Additional finance characteristics (including spatial data) for each project are available from the Chinese Overseas Development Finance (CODF) dataset (Ray et al. 2021).

| ID | Project Description | Country | Year Financed | Project Type | Project Type Details |
|---|---|---|---|---|---|
| 123 | Faleolo International Airport Terminal | Samoa | 2015 | Airport | |
| 396 | Maurice Bishop International Airport Upgrade | Grenada | 2017 | Airport | |
| 792 | Roberts International Airport Rehabilitation | Liberia | 2017 | Airport | |
| 991 | Expansion, Upgrading of Ibrahim Nasir International Airport in Hulhule' | Maldives | 2015 | Airport | |
| 993 | Construction and Development of the Seaplane Facilities at VIA | Maldives | 2018 | Airport | Includes land reclamation |
| 565 | Luanda/Soyo Highway Construction (Bridge over M'Bridge River) | Angola | 2011 | Bridge | |
| 262 | Maputo-Catembe Bridge Construction | Mozambique | 2012 | Bridge | |
| 992 | China Maldives Friendship Bridge | Maldives | 2016 | Bridge | |
| 130 | Foundiougne Bridge Construction | Senegal | 2016 | Bridge | |
| 462 | Karnaphuli River Bridge | Bangladesh | 2014 | Bridge | |
| 141[a] | Abaco infrastructure works: Little Abaco Bridge | Bahamas | 2012 | Bridge | |
| 20 | Convention Centre and Government Offices | Samoa | 2008 | Facility | Offices |
| 42 | Rehabilitation of Sam Lord's Castle Hotel | Barbados | 2015 | Facility | Hotel |
| 419 | Tonga High school Sports Complex for 2019 Pacific Games | Tonga | 2017 | Facility | Sports complex |
| 801 | Attorney General's Office/Mozambican Office of Auditor | Mozambique | 2011 | Facility | Offices |
| 672 | National Telecom Broadband Network Project (Phase II) | Cameroon | 2015 | Facility | Offices |
| 532 | Bunkering Facility and Tank Farm Project at Hambantota port | Sri Lanka | 2009 | Facility | Port facilities |
| 767 | Cape Coast Kotokuraba Market Project | Ghana | 2012 | Facility | Marketplace |
| 656 | Limbe Stadium Construction | Cameroon | 2009 | Facility | Stadium |
| 994 | Development of 1000 Housing Units in Hulhumale | Maldives | 2010 | Facility | Housing development |
| 997 | Development of 1530 Housing Units in Hulhumale | Maldives | 2017 | Facility | Housing development |
| 1014 | West Side of the Lae Tidal Basin and Huon Industrial Park | Papua New Guinea | 2016 | Facility | Industrial Park |
| 544 | Oil refinery | Venezuela | 2010 | Oil Refinery | |
| 229 | Sino-Myanmar Pipeline | Myanmar | 2009 | Pipeline | |
| 805 | Beira Fishing Port Rehabilitation | Mozambique | 2014 | Port | Fishing port |
| 3 | Abidjan Port Expansion | Cote d'Ivoire | 2014 | Port | Shipping port |
| 141[a] | Abaco infrastructure works: North Abaco Port | Bahamas | 2012 | Port | Shipping port |
| 225 | Kribi Port Project (Phase I) | Cameroon | 2011 | Port | Shipping port |
| 311 | Nouakchott Friendship Port Expansion | Mauritania | 2009 | Port | Shipping port |
| 381 | Santiago de Cuba port | Cuba | 2015 | Port | Shipping port |
| 452 | St John Port | Antigua & Barbuda | 2016 | Port | Shipping port |
| 703 | Kribi Port Project (Phase II); PEBC portion | Cameroon | 2016 | Port | Shipping port |
| 707 | Doraleh Multipurpose Port and Damerjob Livestock Port Project | Djibouti | 2016 | Port | Shipping port |

| | | | | | |
|---|---|---|---|---|---|
| 952 | Hambantota Deep Sea Port Phase II (PPP) | Sri Lanka | 2012 | Port | Shipping port |
| 592 | Cabinda Breakwater Construction | Angola | 2016 | Port | Shipping port |
| 33 | PLTU Nangroe Aceh Darussalam Thermal Power Plant | Indonesia | 2009 | Power Plant | Coal |
| 38 | Tanjung Kasam Power Station with Sinosure | Indonesia | 2014 | Power Plant | Coal |
| 45 | Bengkulu Power Station | Indonesia | 2016 | Power Plant | Coal |
| 52 | Paiton Power Plant Unit 9 | Indonesia | 2008 | Power Plant | Coal |
| 53 | Vinh Tan 2 Power Plant | Vietnam | 2010 | Power Plant | Coal |
| 55 | Vinh Tan 1 Coal-Fired Thermal Power Plant | Vietnam | 2014 | Power Plant | Coal |
| 61 | Teluk Sirih Coal-Fired Power Plant (224MW) | Indonesia | 2013 | Power Plant | Coal |
| 63 | Adipala Power Station | Indonesia | 2009 | Power Plant | Coal |
| 70 | Celukan Bawang | Indonesia | 2012 | Power Plant | Coal |
| 86 | Cilacap Sumber Power Station with BOC and Bank Rakyat Indonesia | Indonesia | 2013 | Power Plant | Coal |
| 154 | Hubco Coal Power Plant | Pakistan | 2017 | Power Plant | Coal |
| 171 | Parit Baru Power Station | Indonesia | 2011 | Power Plant | Coal |
| 174 | Payra 1320 MW Thermal Power Plant Project (Kalapara Phase I) | Bangladesh | 2016 | Power Plant | Coal |
| 228 | Vung Ang Power Station | Vietnam | 2011 | Power Plant | Coal |
| 309 | Norochcholai (Lakvijaya) Power Plant Phase 2 | Sri Lanka | 2009 | Power Plant | Coal |
| 336 | Pelabuhan Ratu Power Station | Indonesia | 2009 | Power Plant | Coal |
| 354 | Takalar Steam Coal-Fired Power Plant (200MW) | Indonesia | 2014 | Power Plant | Coal |
| 359 | Quang Ninh-1 Unit 1 | Vietnam | 2009 | Power Plant | Coal |
| 367 | Rembang Power Station | Indonesia | 2008 | Power Plant | Coal |
| 398 | Pacitan Coal Power Plant | Indonesia | 2009 | Power Plant | Coal |
| 403 | Hai Phong Thermal Power Plant Phase 2 | Vietnam | 2008 | Power Plant | Coal |
| 408 | Java-7 Coal-Fired Power Plant (2000MW) | Indonesia | 2016 | Power Plant | Coal |
| 434 | Indramayu Sumuradem Power Station | Indonesia | 2008 | Power Plant | Coal |
| 495 | Cilacap Power Plant Extension Project | Indonesia | 2013 | Power Plant | Coal |
| 935 | Nagan Raya Thermal Power Plant (Meulaboh Power Station) | Indonesia | 2009 | Power Plant | Coal |
| 957 | Vinh Tan Coal Fired Power Plant III Unit I, II, III | Vietnam | 2015 | Power Plant | Coal |
| 111 | Duyen Hai 1 with Sinosure | Vietnam | 2011 | Power Plant | Coal |
| 112 | Duyen Hai 3 with Sinosure/BOC & ICBC | Vietnam | 2012 | Power Plant | Coal |
| 113 | Duyen Hai 2 Thermal Power Plant | Vietnam | 2017 | Power Plant | Coal |
| 334 | Pangkalan Susu Unit 3 & 4 Coal Fired Power Plant | Indonesia | 2013 | Power Plant | Coal |
| 931 | Pangkalan Susu Power Plant Phase II Unit I | Indonesia | 2014 | Power Plant | Coal |
| 1040 | Hunutlu Thermal Power Plant Project with ICBC, BOC | Turkey | 2019 | Power Plant | Coal |
| 526 | Mabini LNG hub | Philippines | 2019 | Power Plant | Gas |
| 332 | Karachi Nuclear Power Complex (K-2/K-3) | Pakistan | 2014 | Power Plant | Nuclear |
| 743 | Hirgigo Thermal Power Plant Upgrade | Eritrea | 2014 | Power Plant | Oil |
| 996 | STELCO 5th Power Development | Maldives | 2016 | Power Plant | Oil |
| 91 | East Coast Rail Link | Malaysia | 2016 | Railway | |

| | | | | |
|---|---|---|---|---|
| 277 | SGR Phase I - Mombasa to Nairobi (Commercial Loan portion) | Kenya | 2014 | Railway |
| 517 | Expansion And Reconstruction Of Existing Line ML-1 | Pakistan | 2017 | Railway |
| 816 | Lagos-Ibadan Railway Modernization (Project II) | Nigeria | 2017 | Railway |
| 104 | Dhaka-Chittagong railway | Bangladesh | 2014 | Railway |
| 156 | Subic-Clark Railway Project | Philippines | 2016 | Railway |
| 959 | National Infrastructure Projects (Dalian IV) | Suriname | 2016 | Road |
| 955 | Colombo-Katunayake Expressway (Construction) | Sri Lanka | 2008 | Road |
| 956 | E01 Southern Expressway (Construction) | Sri Lanka | 2015 | Road |
| 5 | Abidjan-Grand Bassam Highway Construction | Cote d'Ivoire | 2012 | Road |
| 197 | Karachi-Lahore highway | Pakistan | 2014 | Road |
| 203 | North/South toll highway construction | Jamaica | 2013 | Road |
| 493 | East Coast Demerara highway | Guyana | 2017 | Road |
| 564 | Luanda/Soyo Highway Construction (for Nzeto/Soyo section) | Angola | 2011 | Road |
| 566 | Luanda/Soyo Highway Construction (Package 5) | Angola | 2011 | Road |
| 567 | Luanda/Soyo Highway Construction (Package 6) | Angola | 2011 | Road |
| 683 | Yaoundé-Douala Highway Construction (Phase I) | Cameroon | 2012 | Road |
| 44 | Construction of Bar-Boljare Motorway | Montenegro | 2014 | Road |
| 949 | Peshawar-Karachi Motorway (PKM) Project | Pakistan | 2016 | Road |
| 93 | Manado-Bitung Toll Road | Indonesia | 2017 | Road |
| 48 | Road Upgrading Project Nabouwalu Dreketi | Fiji | 2012 | Road |
| 127 | National Road No.3 Construction Project from Phnom Penh (Chom Chao)-Bek Kus-Kampot Town | Cambodia | 2018 | Road |
| 230 | Buca Bay and Moto Road improvement | Fiji | 2013 | Road |
| 231 | Road Reconstruction | Vanuatu | 2014 | Road |
| 313 | National Road Improvement Program | Tonga | 2010 | Road |
| 343 | Benguela, EN100 Road Rehabilitation, Cabo Ledo-Lobito (Lot 5), Ponte do Rio Eval/Ponte do Rio Culango | Angola | 2016 | Road |
| 344 | Cuanza-Sul, EN100 Rehabilitation, Cabo Ledo-Lobito Road (Lot 4), Sumbe-Ponte do Rio Eval | Angola | 2016 | Road |
| 345 | Cuanza-Sul, EN100 Road Rehabilitation, Cabo Ledo-Lobito Road (Lot 2), Ponte do Rio Longa-Ponte do Rio Keve | Angola | 2016 | Road |
| 355 | Rehabiitation & Improvement of Puttalam-Marichchikade-Mannar Road | Sri Lanka | 2011 | Road |
| 386 | Roads Improvement Sigatoka-Serea | Fiji | 2014 | Road |
| 606 | Cuanza-Sul EN100 Road Rehabilitation | Angola | 2016 | Road |
| 766 | Road Construction (Farasol Mbega to Port-Gentil) | Gabon | 2016 | Road |
| 930 | Balikpapan – Samarinda Road Development | Indonesia | 2010 | Road |
| 249 | EN6 Road Repair (Beira to Machipanda); 287km (Loan 1) | Mozambique | 2013 | Road |
| 296 | Nassau Airport Gateway Project | Bahamas | 2011 | Road |
| 934 | Solo-Kertosono Toll Road Project | Indonesia | 2014 | Road |

| | | | | |
|---|---|---|---|---|
| 164 | Luanda-Soyo Highway; Nzeto-Soyo section (Package 7) | Angola | 2011 | Road |
| 162 | Rehabilitation and Improvement of Navatkuli-Karaitivu-Mannar Road | Sri Lanka | 2011 | Road |
| 645 | Takoradi Port Expansion (Phase I) - Access Road | Ghana | 2011 | Road |
| 661 | South Atlantic Inter Link (SAIL) Project | Cameroon | 2015 | Submarine Cable |
| 1009 | Kumul Submarine Cable | Papua New Guinea | 2013 | Submarine Cable |
| 582 | Soyo-Kapary Transmission and Transformation Project | Angola | 2013 | Transmission Line |
| 857 | Dakar Loop Power Transmission (Phase II) | Senegal | 2010 | Transmission Line |

[a] Single loan split into two distinct projects.

**Supplementary Table 2.** Specifications of taxonomic groups considered for analysis.

| Taxonomic Group | Kingdom | Phylum | Class | Subclass |
|---|---|---|---|---|
| Fish | Animalia | Chordata | Actinopterygii | |
| Sea birds | Animalia | Chordata | Aves | |
| Elasmobranchs | Animalia | Chordata | Chondrichthyes | Elasmobranchii |
| Mammals | Animalia | Chordata | Mammalia | |
| Reptiles | Animalia | Chordata | Reptilia | |

**Supplementary Table 3.** IUCN Red List search criteria for marine species included in analysis.

| Field | Criteria |
|---|---|
| Taxonomy | KINGDOM: Animalia<br>PHYLUM: Chordata<br>CLASS: Actinopterygii<br>CLASS: Aves<br>CLASS: Reptilia<br>CLASS: Mammalia<br>CLASS: Chondrichthyes<br>ORDER: Carcharhiniformes<br>ORDER: Heterodontiformes<br>ORDER: Hexanchiformes<br>ORDER: Lamniformes<br>ORDER: Orectolobiformes<br>ORDER: Pristiophoriformes<br>ORDER: Rajiformes<br>ORDER: Squaliformes<br>ORDER: Squatiniformes<br>ORDER: Torpediniformes |
| Red List Category | CR – Critically Endangered<br>EN – Endangered<br>VU – Vulnerable |
| Systems | Marine<br>Terrestrial and Marine<br>Freshwater (=Inland waters) and Marine<br>Terrestrial and Freshwater (=Inland waters) and Marine |
| Country Legends | Extant (resident)<br>Extant & Reintroduced |
| Include | Species |

**Supplementary Table 4.** Threatened species ranges included in the analysis. CR = critically endangered; EN = endangered; VU = vulnerable.

| Taxon group | Threatened species within study areas (Red List Category) |
| --- | --- |
| Elasmobranchs (N = 126) | Alopias pelagicus (EN), Alopias superciliosus (VU), Alopias vulpinus (VU), Carcharhinus acronotus (EN), Carcharhinus albimarginatus (VU), Carcharhinus amblyrhynchos (EN), Carcharhinus borneensis (EN), Carcharhinus brachyurus (VU), Carcharhinus brevipinna (VU), Carcharhinus cerdale (CR), Carcharhinus dussumieri (EN), Carcharhinus falciformis (VU), Carcharhinus longimanus (CR), Carcharhinus melanopterus (VU), Carcharhinus obscurus (EN), Carcharhinus perezi (EN), Carcharhinus plumbeus (VU), Carcharhinus porosus (CR), Carcharhinus signatus (EN), Carcharhinus tjutjot (VU), Carcharias taurus (VU), Carcharodon carcharias (VU), Centrophorus atromarginatus (CR), Centrophorus granulosus (EN), Centrophorus isodon (EN), Centrophorus longipinnis (EN), Centrophorus moluccensis (VU), Centrophorus squamosus (EN), Centrophorus tessellatus (EN), Centrophorus uyato (EN), Centroscymnus owstonii (VU), Cephaloscyllium fasciatum (CR), Cephaloscyllium silasi (CR), Cetorhinus maximus (EN), Chaenogaleus macrostoma (VU), Chiloscyllium burmensis (VU), Chiloscyllium griseum (VU), Chiloscyllium hasselti (EN), Chiloscyllium indicum (VU), Dalatias licha (VU), Deania quadrispinosa (VU), Diplobatis guamachensis (VU), Diplobatis picta (VU), Dipturus batis (CR), Echinorhinus brucus (EN), Eusphyra blochii (EN), Galeorhinus galeus (CR), Ginglymostoma cirratum (VU), Ginglymostoma unami (EN), Glyphis gangeticus (CR), Glyphis garricki (CR), Glyphis glyphis (EN), Halaelurus natalensis (VU), Hemigaleus microstoma (VU), Hemipristis elongata (VU), Hemiscyllium hallstromi (VU), Hemiscyllium michaeli (VU), Hemiscyllium strahani (VU), Hemitriakis leucoperiptera (EN), Holohalaelurus favus (EN), Holohalaelurus punctatus (EN), Isogomphodon oxyrhynchus (CR), Isurus oxyrinchus (EN), Isurus paucus (EN), Lamiopsis temminckii (EN), Lamna nasus (VU), Leucoraja circularis (EN), Leucoraja fullonica (VU), Leucoraja wallacei (VU), Mustelus dorsalis (VU), Mustelus griseus (EN), Mustelus higmani (EN), Mustelus manazo (EN), Mustelus minicanis (EN), Mustelus mustelus (VU), Mustelus punctulatus (VU), Narcine brevilabiata (VU), Narcine entemedor (VU), Narcine lingula (VU), Narcine maculata (VU), Nasolamia velox (EN), Nebrius ferrugineus (VU), Negaprion acutidens (VU), Negaprion brevirostris (VU), Notorynchus cepedianus (VU), Odontaspis ferox (VU), Okamejei boesemani (VU), Okamejei hollandi (VU), Oxynotus centrina (VU), Oxynotus japonicus (VU), Paragaleus leucolomatus (VU), Paragaleus tengi (EN), Platyrhina sinensis (EN), Platyrhina tangi (VU), Proscyllium habereri (VU), Pseudoginglymostoma brevicaudatum (CR), Raja radula (EN), Raja undulata (EN), Rhincodon typus (EN), Rhizoprionodon acutus (VU), Rhizoprionodon lalandii (VU), Rhizoprionodon longurio (VU), Rhizoprionodon porosus (VU), Rostroraja alba (EN), Rostroraja equatorialis (VU), Scymnodon ringens (VU), Somniosus microcephalus (VU), Sphyrna corona (CR), Sphyrna lewini (CR), Sphyrna media (CR), Sphyrna mokarran (CR), Sphyrna tiburo (EN), Sphyrna tudes (CR), Sphyrna zygaena (VU), Squalus acanthias (VU), Squalus hemipinnis (VU), Squalus mitsukurii (EN), Squalus montalbani (VU), Squatina aculeata (CR), Squatina armata (CR), Squatina oculata (CR), Squatina squatina (CR), Squatina tergocellatoides (EN), Stegostoma tigrinum (EN), Temera hardwickii (VU), Triaenodon obesus (VU) |
| Fish (N = 101) | Acanthopagrus vagus (VU), Acentrogobius griseus (VU), Albula glossodonta (VU), Amblyglyphidodon batunai (VU), Amblyglyphidodon ternatensis (VU), Anguilla anguilla (CR), Anguilla borneensis (VU), Anguilla japonica (EN), Anguilla luzonensis (VU), Anguilla rostrata (EN), Argyrosomus japonicus (EN), Argyrosomus thorpei (EN), Balistes capriscus (VU), Balistes punctatus (VU), Bathygobius burtoni (EN), Bolbometopon muricatum (VU), Chaetodontoplus vanderloosi (EN), Coilia mystus (EN), Corcyrogobius lubbocki (VU), Coryphopterus alloides (VU), Coryphopterus eidolon (VU), Coryphopterus hyalinus (VU), Coryphopterus lipernes (VU), Coryphopterus personatus (VU), Coryphopterus thrix (VU), Coryphopterus tortugae (VU), Coryphopterus venezuelae (VU), Cynoglossus macrostomus (VU), Cynoscion acoupa (VU), Dentex dentex (VU), Didogobius amicuscaridis (VU), Elacatinus atronasus (EN), Elacatinus prochilos (VU), Epinephelus akaara (EN), Epinephelus albomarginatus (VU), Epinephelus fuscoguttatus (VU), Epinephelus itajara (VU), Epinephelus marginatus (VU), Epinephelus morio (VU), Epinephelus polyphekadion (VU), Epinephelus striatus (CR), Evynnis cardinalis (EN), Gobiodon aoyagii (VU), Gobiodon axillaris (VU), Gobiodon erythrospilus (VU), Gobiodon fulvus (VU), Gobiodon reticulatus (VU), Gorogobius stevcici (VU), Hippocampus algiricus (VU), Hippocampus barbouri (VU), Hippocampus comes (VU), |

| | |
|---|---|
| | Hippocampus erectus (VU), Hippocampus histrix (VU), Hippocampus ingens (VU), Hippocampus kelloggi (VU), Hippocampus spinosissimus (VU), Hippocampus trimaculatus (VU), Horadandia atukorali (VU), Hyporthodus acanthistius (VU), Hyporthodus flavolimbatus (VU), Hyporthodus niveatus (VU), Kajikia albida (VU), Labrus viridis (VU), Lachnolaimus maximus (VU), Lethrinus mahsena (EN), Lopholatilus chamaeleonticeps (EN), Lucifuga lucayana (EN), Lucifuga spelaeotes (VU), Lutjanus cyanopterus (VU), Makaira nigricans (VU), Meiacanthus abruptus (VU), Merluccius senegalensis (EN), Mola mola (VU), Mycteroperca interstitialis (VU), Nemipterus virgatus (VU), Omobranchus hikkaduwensis (VU), Omobranchus mekranensis (VU), Omobranchus smithi (VU), Oxymonacanthus halli (VU), Oxymonacanthus longirostris (VU), Parablennius lodosus (VU), Paraclinus fehlmanni (VU), Pentanemus quinquarius (VU), Plectropomus areolatus (VU), Plectropomus marisrubri (VU), Polysteganus praeorbitalis (VU), Pomatomus saltatrix (VU), Pseudotolithus senegalensis (EN), Pseudotolithus senegallus (VU), Pseudupeneus prayensis (VU), Rhomboplites aurorubens (VU), Sardinella maderensis (VU), Scarus trispinosus (EN), Sciades parkeri (VU), Siganus niger (VU), Stiphodon rubromaculatus (CR), Thunnus obesus (VU), Thunnus thynnus (EN), Trachurus indicus (VU), Trachurus trachurus (VU), Umbrina cirrosa (VU) |
| Mammals (N = 19) | Aonyx cinereus (VU), Balaenoptera borealis (EN), Balaenoptera musculus (EN), Balaenoptera physalus (VU), Dugong dugon (VU), Eubalaena glacialis (CR), Hippopotamus amphibius (VU), Lutra sumatrana (EN), Lutrogale perspicillata (VU), Monachus monachus (EN), Neophocaena phocaenoides (VU), Orcaella brevirostris (EN), Physeter macrocephalus (VU), Sousa chinensis (VU), Sousa plumbea (EN), Sousa sahulensis (VU), Sousa teuszii (CR), Trichechus manatus (VU), Trichechus senegalensis (VU) |
| Reptiles (N = 14) | Aipysurus fuscus (EN), Batagur affinis (CR), Batagur baska (CR), Batagur borneoensis (CR), Caretta caretta (VU), Carettochelys insculpta (EN), Chelonia mydas (EN), Crocodylus acutus (VU), Dermochelys coriacea (VU), Eretmochelys imbricata (CR), Lepidochelys olivacea (VU), Pelochelys bibroni (VU), Pelochelys signifera (VU), Trionyx triunguis (VU) |
| Sea birds (N = 64) | Anas luzonica (VU), Ardenna bulleri (VU), Ardenna creatopus (VU), Ardeola idae (EN), Aythya ferina (VU), Calidris pygmaea (CR), Calidris tenuirostris (EN), Diomedea dabbenena (CR), Egretta eulophotes (VU), Eurostopodus nigripennis (VU), Falco cherrug (EN), Fregata andrewsi (CR), Fregata aquila (VU), Glareola ocularis (VU), Haliaeetus sanfordi (VU), Hydrobates leucorhous (VU), Hydrobates matsudairae (VU), Larus audouinii (VU), Leptoptilos javanicus (VU), Macrocephalon maleo (EN), Marmaronetta angustirostris (VU), Morus capensis (EN), Mycteria cinerea (EN), Nesofregetta fuliginosa (EN), Numenius madagascariensis (EN), Numenius tenuirostris (CR), Phalacrocorax capensis (EN), Phalacrocorax nigrogularis (VU), Phoebastria irrorata (CR), Platalea minor (EN), Podiceps auritus (VU), Procellaria aequinoctialis (VU), Procellaria parkinsoni (VU), Pseudobulweria becki (CR), Pseudobulweria macgillivrayi (CR), Pterodroma alba (EN), Pterodroma arminjoniana (VU), Pterodroma axillaris (VU), Pterodroma brevipes (VU), Pterodroma cahow (EN), Pterodroma cervicalis (VU), Pterodroma cookii (VU), Pterodroma deserta (VU), Pterodroma hasitata (EN), Pterodroma leucoptera (VU), Pterodroma madeira (EN), Pterodroma phaeopygia (CR), Pterodroma pycrofti (VU), Pterodroma solandri (VU), Puffinus heinrothi (VU), Puffinus mauretanicus (CR), Puffinus yelkouan (VU), Rissa tridactyla (VU), Saundersilarus saundersi (VU), Spheniscus demersus (EN), Sterna aurantia (VU), Sternula balaenarum (VU), Sternula lorata (EN), Thalassarche carteri (EN), Thalassarche chlororhynchos (EN), Thalassarche eremita (VU), Thalassarche salvini (VU), Tringa guttifer (EN), Vultur gryphus (VU) |

**Supplementary Table 5.** Taxa vulnerability weights for each stressor adapted from Butt et al. (2021). Values represent the expected mean vulnerability for all species within each taxonomic group.

| **Taxon group** | **Taxa stressor** | | | | | | | | | | | | | | |
|---|---|---|---|---|---|---|---|---|---|---|---|---|---|---|---|
| | Habitat loss | Noise pollution | Light pollution | Thermal pollution | Inorganic pollution | Nutrient pollution | Organic pollution | Sediment. | Plastic pollution | Invasive species | Wildlife injury | Biomass removal | Bycatch | Entangle. | Poisons, toxins |
| Elasmobranchs | 0.38697 | 0.03655 | 0.00157 | 0.00836 | 0.19369 | 0.21362 | 0.28757 | 0.20003 | 0.22607 | 0.00227 | 0.02369 | 0.88223 | 0.59432 | 0.28697 | 0.27425 |
| Fish | 0.32936 | 0.09167 | 0.16950 | 0.34268 | 0.39997 | 0.33239 | 0.37642 | 0.30527 | 0.33357 | 0.00000 | 0.04130 | 0.62048 | 0.34440 | 0.14911 | 0.31877 |
| Mammals | 0.40230 | 0.33778 | 0.14207 | 0.12205 | 0.18922 | 0.10748 | 0.32940 | 0.09540 | 0.06595 | 0.15629 | 0.61399 | 0.74095 | 0.74095 | 0.58892 | 0.04637 |
| Reptiles | 0.40655 | 0.00544 | 0.12459 | 0.10135 | 0.24670 | 0.10177 | 0.30583 | 0.23065 | 0.06392 | 0.17616 | 0.53493 | 0.69705 | 0.68951 | 0.54466 | 0.04600 |
| Sea birds | 0.41917 | 0.37790 | 0.48457 | 0.08377 | 0.19338 | 0.07575 | 0.26697 | 0.21691 | 0.05503 | 0.51065 | 0.46748 | 0.68188 | 0.60203 | 0.36523 | 0.03836 |

**Supplementary Table 6.** Habitat vulnerability weights for each stressor, adapted from Halpern et al. (2008) and rescaled between 0 and 1.

| Habitat | | Habitat stressor | | | | | | | | | | | |
|---|---|---|---|---|---|---|---|---|---|---|---|---|---|
| | | Light pollution | Sea surface temp. | Inorganic pollution | Nutrient pollution | Ocean pollution | Direct human | Benthic structures | Shipping | Invasive species | Demersal, destructive fishing | Commercial fishing[a] | Artisanal fishing |
| Intertidal | Rocky intertidal | 0.28 | 0.56 | 0.42 | 0.32 | 0.26 | 0.56 | 0.20 | 0.06 | 0.56 | 0.24 | 0.12 | 0.26 |
| | Mud flats | 0.28 | 0.28 | 0.32 | 0.32 | 0.16 | 0.44 | 0.18 | 0.38 | 0.58 | 0.28 | 0.16 | 0.08 |
| | Beach | 0.40 | 0.12 | 0.12 | 0.08 | 0.10 | 0.54 | 0.16 | 0.38 | 0.18 | 0.04 | 0.08 | 0.14 |
| | Mangroves | 0.18 | 0.48 | 0.10 | 0.36 | 0.24 | 0.66 | 0.26 | 0.40 | 0.20 | 0.00 | 0.08 | 0.34 |
| | Salt marsh | 0.36 | 0.28 | 0.40 | 0.38 | 0.24 | 0.32 | 0.18 | 0.28 | 0.56 | 0.20 | 0.14 | 0.12 |
| Coastal | Coral reefs | 0.20 | 0.56 | 0.14 | 0.36 | 0.24 | 0.46 | 0.10 | 0.30 | 0.30 | 0.24 | 0.20 | 0.46 |
| | Seagrass beds | 0.10 | 0.42 | 0.16 | 0.42 | 0.10 | 0.50 | 0.32 | 0.38 | 0.24 | 0.04 | 0.10 | 0.06 |
| | Kelp forests | 0.10 | 0.40 | 0.00 | 0.08 | 0.02 | 0.32 | 0.00 | 0.00 | 0.26 | 0.30 | 0.18 | 0.16 |
| | Rocky reefs | 0.14 | 0.38 | 0.44 | 0.32 | 0.34 | 0.50 | 0.34 | 0.28 | 0.50 | 0.54 | 0.52 | 0.44 |
| | Shellfish reefs | 0.20 | 0.16 | 0.54 | 0.28 | 0.00 | 0.60 | 0.08 | 0.00 | 0.52 | 0.62 | 0.06 | 0.20 |
| | Shallow, soft bottom | 0.10 | 0.10 | 0.30 | 0.40 | 0.22 | 0.40 | 0.02 | 0.06 | 0.54 | 0.42 | 0.22 | 0.00 |
| Offshore | Continental shelf, soft bottom | 0.00 | 0.50 | 0.42 | 0.28 | 0.24 | 0.22 | 0.10 | 0.34 | 0.32 | 0.60 | 0.28 | 0.18 |
| | Continental shelf, hard bottom | 0.00 | 0.58 | 0.04 | 0.34 | 0.06 | 0.58 | 0.42 | 0.18 | 0.30 | 0.62 | 0.56 | 0.38 |
| | Continental slope, soft bottom | 0.00 | 0.46 | 0.42 | 0.40 | 0.28 | 0.00 | 0.32 | 0.02 | 0.04 | 0.64 | 0.22 | 0.00 |
| | Continental slope, hard bottom | 0.00 | 0.18 | 0.04 | 0.12 | 0.34 | 0.00 | 0.44 | 0.20 | 0.10 | 0.56 | 0.22 | 0.08 |
| | Deep benthic, soft bottom | 0.00 | 0.50 | 0.36 | 0.26 | 0.46 | 0.32 | 0.38 | 0.18 | 0.30 | 0.46 | 0.28 | 0.06 |
| | Deep benthic, hard bottom | 0.00 | 0.30 | 0.00 | 0.00 | 0.24 | 0.00 | 0.32 | 0.00 | 0.00 | 0.60 | 0.00 | 0.00 |
| | Seamounts | 0.00 | 0.36 | 0.00 | 0.00 | 0.24 | 0.00 | 0.28 | 0.00 | 0.00 | 0.70 | 0.00 | 0.18 |
| | Shallow pelagic | 0.08 | 0.66 | 0.46 | 0.24 | 0.34 | 0.18 | 0.30 | 0.38 | 0.46 | 0.42 | 0.40 | 0.20 |
| | Deep pelagic | 0.00 | 0.46 | 0.32 | 0.00 | 0.08 | 0.00 | 0.00 | 0.00 | 0.00 | 0.16 | 0.14 | 0.00 |

[a] Represents the mean vulnerability across four stressors: demersal non-destructive fishing (high and low bycatch) and pelagic non-destructive fishing (high and low bycatch)

**Supplementary Table 7.** Links between taxa stressors identified by Butt et al. (2021) and habitat stressors identified by Halpern et al. (2008) used to calculate exposure risks to habitats. Habitat stressors reflecting multiple taxa-based stressors (i.e. direct human, shipping, and fishing stressors) are assumed to reflect the average of each associated taxa stressor. Some stressors (e.g. thermal pollution and sea surface temperature) are not directly comparable, but are assumed to threaten marine systems through similar physiological mechanisms.

| | **Habitat stressor** | | | | | | | | | | | | | | |
|---|---|---|---|---|---|---|---|---|---|---|---|---|---|---|---|
| **Taxa stressor** | Light pollution | Sea surface temp. | Inorganic pollution | Nutrient pollution | Ocean pollution | Direct human | Benthic structures[a] | Shipping | Invasive species | Demersal, destructive fishing | Demersal fishing, high bycatch[b] | Demersal fishing, low bycatch[b] | Pelagic fishing, high bycatch[b] | Pelagic fishing, low bycatch[b] | Artisanal fishing |
| Habitat loss | | | | | | **X**[d] | **X** | | | **X**[e] | | | | | |
| Noise pollution | | | | | | **X**[d] | | **X**[d] | | | | | | | |
| Light pollution | **X** | | | | | | | | | | | | | | |
| Thermal pollution | | **X** | | | | | | | | | | | | | |
| Inorganic pollution | | | **X** | | | | | | | | | | | | |
| Nutrient pollution | | | | **X** | | | | | | | | | | | |
| Organic pollution | | | | | **X** | | | | | | | | | | |
| Sedimentation[c] | | | | | | | | | | | | | | | |
| Plastic pollution | | | | | | **X**[d] | | **X**[d] | | | | | | | **X**[f] |
| Invasive species | | | | | | | | | **X** | | | | | | |
| Wildlife injury | | | | | | | | **X**[d] | | | | | | | |
| Biomass removal | | | | | | | | | | **X**[e] | **X**[d] | **X**[d] | **X**[d] | **X**[d] | **X**[f] |
| Bycatch | | | | | | | | | | **X**[e] | **X**[d] | **X**[d] | **X**[d] | **X**[d] | **X**[f] |
| Entanglement | | | | | | | | | | **X**[e] | **X**[d] | **X**[d] | **X**[d] | **X**[d] | **X**[f] |
| Poisons, toxins | | | | | | | | | | | | | | | **X**[f] |

[a] Specific to pipelines and submarine cables
[b] Stressors are aggregated in the analysis as "Commercial fishing"
[c] No equivalent habitat stressor (excluded from impact risk estimates to habitats)
[d] Exposure risk divided by 3
[e] Exposure risk divided by 4
[f] Exposure risk divided by 5
